# *Epichloë festucae* in mutualistic association with *Lolium perenne* suppresses host apoplastic cysteine protease activity

**DOI:** 10.1101/2020.11.06.371211

**Authors:** Andrea Passarge, Fatih Demir, Kimberly Green, Jasper R.L. Depotter, Barry Scott, Pitter F. Huesgen, Gunther Doehlemann, Johana C. Misas Villamil

**Affiliations:** Institute for Plant Sciences, University of Cologne, Cologne, Germany; Central Institute for Engineering, Electronics and Analytics, Forschungszentrum Jülich, Jülich, Germany; School of Fundamental Sciences, Massey University, Palmerston North, New Zealand; Bio-Protection Research Centre, Massey University, Palmerston North, NZ; Cologne Excellence Cluster for Stress Responses in Ageing-Associated Diseases (CECAD), University of Cologne, Cologne, Germany; Institute for Biochemistry, University of Cologne, Cologne, Germany

**Keywords:** apoplast, cystatin, endophyte, papain-like cysteine protease, ryegrass

## Abstract

Plants secrete various defence-related proteins into the apoplast, including proteases. Papain-like cysteine proteases (PLCPs) are central components of the plant immune system. To overcome plant immunity and successfully colonise their hosts, several plant pathogens secrete effector proteins inhibiting plant PLCPs. We hypothesized that not only pathogens but also mutualistic microorganisms interfere with PLCP-meditated plant defences to maintain endophytic colonisation with their hosts. *Epichloë festucae* forms mutualist associations with cool season grasses and produces a range of secondary metabolites that protect the host against herbivores. In this study, we performed a genome wide identification of *Lolium perenne* PLCPs, analysed their evolutionary relationship and classified them into nine PLCP subfamilies. Using activity-based protein profiling, we identified four active PLCPs in the apoplast of *L. perenne* leaves that are inhibited during endophyte interactions. We characterized the *L. perenne* cystatin LpCys1 for its inhibitory capacity against ryegrass PLCPs. LpCys1 inhibits LpCP2, indicating that LpCys1 might play a role in the suppression of PLCP activity during the interaction with *E. festucae*. However, since the activity of other *L. perenne* PLCPs is not sensitive to LpCys1 we propose that additional inhibitors are involved in the suppression of apoplastic PLCPs during *E. festucae* infection.

## Introduction

Plants are continuously exposed to a great variety of microbes ranging from mutualists to pathogens. The epidermal surface and the apoplast are primary interfaces of plant-microbe interactions. The fungal endophyte *Epichloë festucae* forms symbiotic associations with temperate Festuca and Lolium grass hosts (Leuchtmann *et al.*, 1994). Hyphae reside within the intercellular spaces between host cells and systemically colonise the apoplast within the leaf sheath, leaf blade and inflorescences (May *et al.*, 2008; Scott *et al.*, 2012). In the later stages of host development, hyphae cease growing and become closely attached to the host cell wall by an adhesive matrix and remain metabolically active (Tan *et al.*, 2001; Christensen and Voisey, 2007). The intercellular growth of *E. festucae* within *Lolium perenne* is tightly regulated and the loss of key signalling components leads to a disruption in symbiosis (Tanaka *et al.*, 2006, 2008, 2013; Takemoto *et al.*, 2006, 2011; Eaton *et al.*, 2010; Becker *et al.*, 2015). This tight association between *E. festucae* hyphae and the host cell wall, has been proposed to facilitate endophyte-host crosstalk through the exchange of signalling molecules (Eaton *et al.*, 2011). To successfully colonise the host and survive within the apoplast, it has been proposed that *E. festucae* needs to suppress host defences (Schardl *et al.*, 2004; Scott *et al.*, 2018). The apoplast is a harsh environment that harbours hydrolytic enzymes such as chitinases and proteases, shown to be involved in plant defence response to microbes (Ökmen *et al.*, 2018; Thomas and van der Hoorn, 2018). Among these, apoplastic proteases such as Papain-Like Cysteine Proteases (PLCPs) are hubs in plant immunity (Misas Villamil *et al.*, 2016). PLCPs may release damage or microbe associated molecular patterns (DAMPs or MAMPs) as well as small signalling peptides which activate signalling cascades triggering the induction of defense responses (Ziemann *et al.*, 2018; Paulus *et al.*, 2020). They can also act as co-receptors and decoys to prevent pathogen colonization (Kourelis *et al.*, 2020). Accordingly, distant related plant pathogens have evolved effectors targeting PLCPs or their regulators, highlighting their importance in plant immunity (Krüger *et al.*, 2002; Rooney *et al.*, 2005; Song *et al.*, 2009; Kaschani *et al.*, 2010; Bozkurt *et al.*, 2011; Lozano-Torres *et al.*, 2012; Clark *et al.*, 2018; Misas Villamil *et al.*, 2019). Pathogens can also manipulate the host to produce plant cystatins, cysteine protease inhibitors, to overcome defense responses. One example is CC9, a maize cystatin that suppresses host immunity during *Ustilago maydis* infection by inhibiting PLCPs (van der Linde *et al.*, 2012*a*). It has been proposed that like pathogenic fungi, plant endophytic fungi secrete effector molecules to promote host colonization (Zamioudis and Pieterse, 2012; Spanu and Panstruga, 2017). However, so far only a small number of effectors from mutualistic fungi, mostly mycorrhiza, have been identified and characterized (Kloppholz *et al.*, 2011; Plett *et al.*, 2011, 2014; Wawra *et al.*, 2016; Perotto *et al.*, 2018; Nostadt *et al.*, 2020). Within the *E. festucae* Fl1 genome, 158 small (< 300 amino acid residues in length) secreted protein-encoding genes are predicted (Hassing *et al.*, 2019). Many of these are highly expressed *in planta*, and differentially regulated during pathogenic *E. festucae* associations caused by single gene deletions (Eaton *et al.*, 2010; Schardl *et al.*, 2013).

Mutations in *E. festucae* genes that disrupt cell-cell fusion and other key signalling pathways lead to an antagonistic interaction characterized by unregulated growth of endophytic hyphae and detrimental effects on host growth (Scott *et al.*, 2018). Furthermore, key components of the activation of immune responses such as host pathogenesis related (PR) and respiratory burst responses genes are down regulated during mutualistic *E. festucae* associations (Dupont *et al.*, 2015). Although these findings have made significant contributions towards our understanding of the *E. festucae* - host system, it is still largely unknown how *E. festucae* successfully modulates host defence responses. In this study, we investigate the role of apoplastic PLCPs of *L. perenne* during the interaction with *E. festucae*. Through computational and proteomic approaches, we identified several PLCPs present and active in the leaf apoplast of *L. perenne*. We show that commonly active PLCPs of uninfected plants are inhibited in response to *E. festucae* interaction. We further identified an apoplastic *L. perenne* - derived cystatin and analysed its inhibitory effect on apoplastic PLCPs.

## Materials and methods

### Plant material

*Lolium perenne* cv Samson and *Nicotiana benthamiana* plants were grown in greenhouse with long day period (16 h light) at 23°C and 8 h dark period at 20 °C with 30 – 40% humidity. *Lolium perenne* infected Fl1 and CT plants were kindly provided by Dr. Yvonne Becker (JKI, Julius Kühn - Institute, Braunschweig, Germany).

### Strains and plasmid construction

The Golden gate modular cloning system was used to generate plasmids used for the heterologous expression of PLCPs in *N. benthamiana*. All Oligonucleotides used for PCR are listed in suppl. Table S1. To obtain pL1M-F1-LpCP2::2×35S, pL1M-F1-LpXCP2::2×35S and pL1M-F1-LpCathB::2×35S, LpCP2 (maker-scaffold_4870|ref0016801-exonerate_est2genome-gene-0.3), LpXCP2 (maker-scaffold_182|ref0000331-exonerate_est2genome-gene-0.0) and LpCathB (maker-scaffold_11872|ref0015306-exonerate_est2genome-gene-0.3) were amplified from *L. perenne* cDNA via PCR. pL1M-F1-LpCP1::2×35S was amplified from LpCP1 (maker-scaffold_2516|ref0039699-exonerate_est2genome-gene-0.3) leaving out the DNA sequence coding for the granulin domain. pL1M-F1-CP1Amut-nogran-mCherry::2×35S is described in Schulze Hüynck *et al.*, 2019. The DNA fragments were ligated as previously described in Weber *et al.*, 2011 and transformed first into *E. coli* Top10 competent cells and then into *A. tumefaciens* GV3101 competent cells for overexpression in *N. benthamiana*. For the expression of LpCys1 in *E. coli*, R_12141 (maker-scaffold_12141|ref0031444-exonerate_est2genome-gene-0.0) was amplified without signal peptide using *L. perenne* cDNA. Subsequently, the PCR product was ligated with the PvuII-HF digested plasmid pRSET-GST-PP to obtain pRSET-PP-LpCys1-noSP, which was transformed to *E. coli* BL21 (DE3)pLysS competent cells. All strains used are listed in suppl. Table S2.

### Heterologous expression of PLCPs in N. benthamiana leaves

*Agrobacterium tumefaciens*, containing the desired construct, were grown at 28°C in liquid dYT media, supplemented with the appropriate antibiotics, until an OD_600_ between 0.8 and 1.6 was reached. The cultures were diluted with 10 mM magnesium chloride to a final OD_600_ of 1. After at least one h incubation in darkness with 200 μM acetosyringone (Sigma-Aldrich, Taufkirchen, Germany), 5 - 6 weeks old *N. benthamiana* leaves were infiltrated with a needleless tuberculin-syringe.

### Apoplastic fluid isolation from N. benthamiana and L. perenne leaves

Isolation of *N. benthamiana* apoplastic fluids was performed as described in Schulze Hüynck *et al.*, 2019. In short, three days post *Agrobacterium* infiltration *N. benthamiana* leaves were harvested, and vacuum infiltrated with MilliQ water three times for 5 min at 60 mbar with a 2 min interval of atmospheric pressure. The leaves were surface dried, transferred to Falcon tubes and centrifuged for 20 min at 2000 *g* to isolate the apoplastic fluid. If not used directly, the apoplastic fluid was stored at −20°C. *L. perenne* leaves were cut ca. 3 cm above soil to avoid damage to the shoot apical meristem. The leaves were gently separated, and vacuum infiltrated with MilliQ water three times for 10 min at 60 mbar with a 2 min interval of atmospheric pressure and otherwise treated as *N. benthamiana* leaves.

### Activity Based Protein Profiling (ABPP)

Leaf apoplastic fluid was incubated in darkness for 2 h at room temperature (RT) in 50 mM sodium acetate, 10 mM DTT, DMSO and 0.2 μM of activity based probe MV201 or MV202 (Richau *et al.*, 2012). Prior to labelling one set of samples were pre-incubated with 10 μM or 20 μM E-64 (Sigma-Aldrich, St. louis, Mississippi, USA), as negative control. Labelling was stopped by the addition of 500 μl acetone, followed by protein precipitation overnight at −20°C. The supernatant was discarded after samples were centrifuged for 30 min at max. speed and 4°C. The pellet was resuspended in water and 1 × SDS-loading dye. Samples were boiled for 5 min at 95°C and separated via SDS-PAGE using 12% or 15% SDS gels. MV202 and MV201 labelled proteins were visualised by in-gel fluorescent scanning using a rhodamine filter (Ex. 532 nm, Em. 580 nm) using the Chemi-Doc MP System (Bio-Rad, California, USA). Sample loading was visualised via SyproRuby stain (Ex. 450 nm, Em. 610 nm; SyproRuby Invitrogen, Carlsbad, California, USA), performed according to manufacturer’s instructions. The Quantification of PLCP-signal via rhodamine signal strength was performed using the ImageLabTM software (Bio-Rad, Hercules, CA, United States). For convolution ABPPs, apoplastic fluid of mock and *E. festucae* infected *L. perenne* plants was mixed in a 1:1 ratio and incubated for 1 h at RT prior to labelling. Samples were labelled with MV202 as described above. After labelling, the apoplastic fluid of mock and *E. festucae* infected *L. perenne* plants was mixed in a 1:1 ratio. Subsequently, samples were treated as previously described. For the inhibition assays, apoplastic fluid was extracted from *N. benthamiana* plants expressing *L. perenne* PLCPs. Prior to labelling the apoplastic fluid was incubated for 30 min with different concentrations of recombinant LpCys1 or chicken egg white cystatin (CEWC, Sigma-Aldrich, St. louis, Mississippi, USA) and Tris-HCl (24 mM Tris, 71 mM NaCl; pH 7.5). As negative control one set of samples was pre-incubated with 20 μM E-64 (Sigma-Aldrich, St. louis, Mississippi, USA). Subsequently, samples were treated as described above.

### PLCP pulldown using streptavidin beads

Leaf apoplastic fluid of three uninfected *L. perenne* plants was isolated as described before. Apoplastic fluid (2.35 ml) was incubated for 4 h at RT in 50 mM sodium acetate pH 6, 10 mM DTT and 2 mM DCG-04 (Greenbaum *et al.*, 2000). As a negative control, one set of samples was incubated with an equivalent amount of DMSO instead of DCG-04. The pulldown experiment was subsequently performed as described in Schulze Hüynck *et al.*, 2019.

### Apoplast proteome sample preparation

Three biological replicates of ryegrass apoplast fluid were collected, proteins purified by chloroform/methanol precipitation, cysteine residues reduced with 10 mM DTT and alkylated with 30 mM IAA and digested with MS-grade Trypsin (Serva) for 16 h at 37°C. Stable isotope labelling was achieved by reductive dimethylation of peptide N-terminal and Lys side chain primary amines with 20 mM CH_2_O and 20 mM NaBH_3_CN (+28.0313 Da) for mock treated plants, 20 mM CD_2_O and 20 mM NaBH_3_CN (+32.0564 Da) for CT-infected plants and 20 mM ^13^CD_2_O and 20 mM NaBD_3_CN (+36.0756 Da) for Fl1 treatment (Boersema *et al.*, 2009). Labelling reactions were quenched with final 100 mM Tris-HCl pH 6.8 for 1h at RT, pooled in a 1:1:1 ratio and subsequently separated in three fractions at high pH (10%/15%/20% ACN, 10 mM NH4OH) followed by a final elution at acidic pH (50% ACN, 0.1% formic acid (FA)). The fractions were evaporated to dryness in a vacuum concentrator and reconstituted in 2% can, 0.1% FA prior to LC/MS analysis.

### Nano LC-MS/MS measurements

LC-MS/MS analysis was performed with an UltiMate 3000 RSCL nano-HPLC system (Thermo) online coupled to an Impact II Q-TOF mass spectrometer (Bruker) via a CaptiveSpray ion source boosted with acetonitrile-saturated nitrogen gas stream. Peptides were loaded on a Acclaim PepMap100 C18 trap column (3 μm, 100 Å, 75 μm i.d. x 2 cm, Thermo) and separated on a Acclaim PepMap RSLC C18 column (2 μm, 100 Å, 75 μm i.d. x 50 cm, Thermo) with a 2h elution protocol that included an 80min separation gradient from 5% to 35% solvent B (solvent A: H_2_O + 0.1% FA, solvent B: can, 0.1% FA) at a flow of 300 nL/minute at 60 °C. Line-mode MS spectra were acquired in mass range 200 – 1400 m/z with a Top14 method at 4 Hz sampling rate for MS1 spectra and an intensity-dependent acquisition rate of 5 to 20 Hz for MS2 spectra. The capillary voltage for the CaptiveSpray ion source was 1600V. Collision energies of 7 eV and 9 eV were applied in two equal steps with the ion transfer time set to 61 and 100 μs, respectively, during acquisition of each MS2 spectrum.

### Mass spectrometry data analysis

Peptides were identified by matching spectra against a combination of a custom *Epichloë festucae* database (EfFl1_Proteins_Annotated_2020-05.fasta, containing 7077 sequences, Aug. 2018), a *Lolium perenne* database (lope_proteins.V1.0.fasta, 40068 entries, downloaded 18/03/2019, Byrne *et al.*, 2015) and the sequences of maize and Arabidopsis PLCPs (PLCPs_Ath+Maize.fasta”, 52 entries) using the Andromeda search engine integrated into the MaxQuant software package (version 1.6.0.16) with standard settings (Tyanova *et al.*, 2016). Carbamidomethylation of cysteine (+ 56.0214 Da) was set as a fixed peptide modification. Oxidation of methionine (+ 15.9949 Da) and acetylation of protein N-termini (+ 42.0106 Da) were set as variable modifications. For the apoplast proteome sample, triplex dimethyl isotope labelling with light ((CH_3_)_2_, +28.0313 Da), medium ((CD_2_H)_2_,+32.0564 Da) and heavy (^13^CD_3_)_2_), (+36.0756 Da) dimethyl label at Lys residues and peptide N-termini was additionally considered. The “requantify” option was enabled and false discovery rates (FDR) for peptide sequence matches and protein identifications were set to < 0.01. Only proteins quantified in at least 2 of the 3 biological replicates were used for pairwise comparisons of each of the three conditions. Protein ratios were median-normalized within each replicate before assessing differential expression with a moderated *t*-test using the “limma” package for R (Ritchie *et al.*, 2015). Proteins changing at least 50% in abundance (log_2_ fold change <−0.58 or > 0.58) supported by a moderated *t*-test p-value < 0.05 and were considered significantly changed in abundance.

### Recombinant expression and purification of LpCys1

The Plasmid pRSET-GST-PP-LpCys1-noSP was transformed into *E. coli* BL21 (DE3)pLysS competent cells (Novagen/Merck, Darmstadt, Germany). An overnight culture grown in dYT medium supplemented with 100 μg/ml carbenicillin and 34 μg/ml chloramphenicol was diluted to an OD_600_ of 0.1 with dYT supplemented with 100 μg/ml carbenicillin and grown at 37°C and 200 rpm to an OD_600_ of 0.6. The LpCys1 expression was induced with 1 mM IPTG. After 4 h at 37°C and 200 rpm, the cells were harvested by centrifuging for 30 min at 4°C and 6,000 rpm (JA-10, Beckman Coulter®). The cells were resuspended in 1x PBS (pH 7.3) and 1 μl benzonase (Sigma-Aldrich, St. louis, Mississippi, USA) and protease inhibitor mix was added. The mix was incubated for 30 min at RT, followed by the addition of 5 mM DTT and sonification. The insoluble cell debris was removed via centrifugation for 30 min at 4°C and 20,000 rpm (JA-25.50, Beckman Coulter®). The supernatant was incubated for 1 h with low agitation and with 1.2 ml Glutathione-sepharose (GE-Healthcare, Uppsala, Sweden) that was equilibrated three times with 12 ml cold 1x PBS (pH 7.3). The mix was subsequently applied to a flow-through column and the column was washed three times with 12 ml 1x PBS (pH 7.3) with 5 mM DTT and once with PreScission cleavage buffer (50 mM Tris-HCl pH 7.5, 150 mM NaCl, 1 mM EDTA, 1 mM DTT). PreScission protease mix was added and incubated on the column overnight at 4 °C. The flow-through was collected. The sepharose matrix was washed three times with 1.2 ml PreScission cleavage buffer and the flow-through was collected and all collected flow-through fractions were pooled. The pooled protein sample was treated as describe in Mueller *et al.*, 2013. The HiLoad® 16/600 Superdex® 75 pg column (GE Healthcare, Chicago, Illinois, USA) with Tris-HCl buffer (50 mM Tris-HCl, 150 mM NaCl, 5 mM DTT, pH 7.5) was used for gel filtration and Vivaspin 15 with 5 MWCO (Sartorius Stedim Biotech GmbH, Goettingen, Germany) was used to concentrate the final protein fractions containing LpCys1.

### Identification and phylogenetic analysis of PLCPs and cystatins in L. perenne

Predicted proteins of *Lolium perenne* were obtained from Byrne *et al.*, 2015 and a functional prediction was carried out using InterProScan (v5.32-71.0). Subsequently it was scanned for PLCP associated PFAM number PF00112 and the program SignalP (v4.1) was used to identify those with an N-terminal secretion signal. All original identifiers are listed in suppl. Table S3. The PLCP phylogenetic tree was generated using the identified PLCPs in *L. perenne* as well as 52 maize PLCP sequences from the MEROPS database (B73 line), six maize PLCPs identified in the early golden bantam line and 39 barley PLCP sequences (Díaz-Mendoza *et al.*, 2014; Rawlings *et al.*, 2018; Schulze Hüynck *et al.*, 2019). One PLCP of each PLCP subfamily from *Arabidopsis thaliana* were also included in the tree (Richau *et al.*, 2012), as well as four *A. thaliana* cysteine proteases of the family C13 (legumains) α-VPE (AEC07775.1), β-VPE (OAP12170.1), γ-VPE (OAO96694.1) and δ-VPE (OAP02173.1). Sequences used for the phylogenetic analysis of plant PLCPs can be found in suppl. Dataset S1. For the construction of the tree MAFFT (v7.407), RAxML (v8.2.12) with the PROTGAMMAWAG substitution model were used (Katoh and Standley, 2013; Stamatakis, 2014). The original phylogenetic tree can be found in suppl. Fig. S1. For the plant cystatins a phylogenetic tree was generated by aligning the full-length protein sequences (suppl. Dataset S2) of the identified apoplastic *L. perenne* cystatins LpCys1, LpCys4 and LpCys9 as well as cystatin sequences from *H. vulgare* (HvCPI), *Z. mays* (CC), *O. sativa* (OC) and *A. thaliana* (AtCYS) using MAFFT (v7.407) (Martinez *et al.*, 2009; Martínez *et al.*, 2012; van der Linde *et al.*, 2012*b*; Katoh and Standley, 2013; Stamatakis, 2014). The unrooted radial tree was generated with RAxML (v8.2.12) using the substitution model PROTGAMMAWAG (Stamatakis, 2014). 100 bootstraps were performed, bootstrap values a given in the tree. The trees were visualised with FigTree (v1.4.2, http://tree.bio.ed.ac.uk/software/figtree).

## Results

### Apoplastic PLCP activity is reduced during E. festucae interaction

Inhibition of apoplastic PLCPs has been shown to be crucial for successful infection by a diversity of pathogens (Misas Villamil *et al.*, 2016) but their role in colonization by fungal endophytes is not known. In nature, different *E. festucae* strains exhibit varying degrees of host specificity and host colonisation (Scott *et al.*, 2018). In our analysis we therefore included two *E. festucae* strains, Fl1 and Common Toxic (CT). Fl1 is a commonly used laboratory strain that can be artificially inoculated into the host *L. perenne*, while CT is a naturally occurring endophyte of *L. perenne*. To determine if the leaf apoplastic PLCP activity of *L. perenne* is modulated during the interaction with *E. festucae*, PLCP activity was monitored using activity-based protein profiling (ABPP; Fig. 1). Leaf apoplastic fluid (AF) was extracted from mock and infected plants. One set of plants was infected with the *E. festucae* strain Fl1 (sexual form), the other set of plants was naturally infected seed with *E. festucae* var. *lolii* strain CT (Common toxic, asexual form). AF was labelled with the activity based probe MV201 which contains an epoxide specific E-64-based inhibitor group that covalently and irreversibly binds to the active site of PLCPs (Richau *et al.*, 2012). A fluorescent BODIPY moiety allows the detection of MV201 labelled PLCPs. Uninfected (E-) plants showed the strongest apoplastic PLCP signals at ca. 35 kDa that were out competed showing a decrease in signal intensity when the PLCP specific inhibitor E-64 was added in excess. In comparison to uninfected plants, apoplastic PLCP activity in *E. festucae* Fl1 and CT infected plants was significantly reduced (Fig. 1A). Notably, AFs from plants infected with the Fl1 strain showed a stronger PLCP signal reduction than AFs from CT infected plants. Furthermore, the SyproRuby loading control showed proteome differences between endophyte infected samples. Despite of a high diversity in protein patterns between treatments, a distinct signal intensity for proteins at ca. 35 kDa was observed in Fl1 samples, which was less intense in E− and CT samples (Fig. 1B). These findings indicate that endophyte infection alters the host apoplastic proteome whereas PLCP activity, a key component of plant immunity, is reduced during *E. festucae* interactions, particularly in response to Fl1 interactions.

**Fig. 1.**
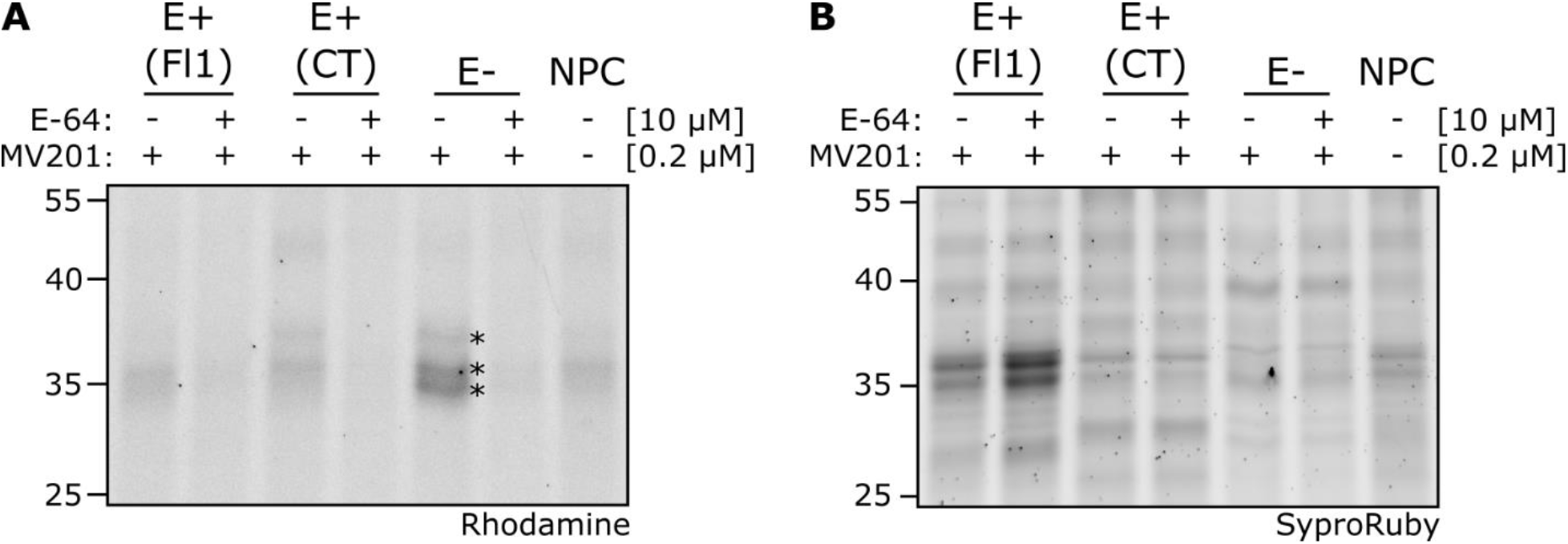
Activity of apoplastic PLCPs in *Epichloe festucae* infected *Lolium perenne* leaves. Leaf apoplastic fluid of endophyte infected (E+) Fl1, CT and mock (E−) plants was isolated. Samples were pre-incubated for 15 min with E-64 or the equivalent amount of DMSO before labelling with the fluorescent activity-based probe MV201. (A) Samples were separated via SDS-PAGE and labelled PLCPs were visualized via in-gel fluorescent scanning using a rhodamine filter (Ex. 532 nm, Em. 580 nm). Asterisks indicate active PLCPs. (B) Sample loading was monitored using SyproRuby staining (Ex. 450 nm, Em. 610 nm). Numbers on the left side of gel pictures indicate the protein ladder in KDa.

To characterize PLCP modulation in *L. perenne* plants it is essential to identify the PLCPs in this plant species and classify them into the nine PLCP subfamilies (Richau *et al.*, 2012). Although PLCPs are crucial for many processes including plant development, senescence and plant defence, the peptidase database MEROPS only lists one PLCP, MER0345289, for *L. perenne*, belonging to the apoplastic PLCP subfamily C1A (Rawlings *et al.*, 2018). In other plant species the number of identified PLCPs ranged from 63, 48, 42, 36, and 30 in maize, sorghum, barley, Arabidopsis and rice, respectively (Díaz-Mendoza *et al.*, 2014; Sekhon *et al.*, 2019). We performed a functional protease domain screen via InterProScan (www.ebi.ac.uk/interpro) using the public genome annotation of Byrne *et al.*, 2015. Subsequently, the presence of PLCP related PFAM domains and IPR identifiers to determine further *L. perenne* PLCPs was evaluated. This search identified 23 *L. perenne* PLCPs containing an N-terminal signal peptide and a C1A protease domain (Suppl. Table S3 and Suppl. Dataset S1). To classify these newly identified PLCPs into the nine PLCP subfamilies (Richau *et al.*, 2012), protein sequences were compared to other plant PLCPs and phylogenetically analysed. The 23 newly identified PLCPs, 58 maize PLCP sequences (Rawlings *et al.*, 2018; Schulze Hüynck *et al.*, 2019), 38 barley sequences (Díaz-Mendoza *et al.*, 2014) and one member of each PLCP subfamily of Arabidopsis (Richau *et al.*, 2012) were used to generate a phylogenetic tree using the maximum likelihood method (Fig. 2). Sequences from the four Arabidopsis cysteine proteases of the family C13 (legumains) α-VPE, β-VPE, γ-VPE and δ-VPE were used as outgroup. This analysis showed that the largest group of *L. perenne* PLCPs belong to the subfamilies THI1 (5 members), SAG12 (5 members) and RD21 (4 members). The remaining seven PLCPs are distributed among the XCP2, RD19A, CTB3, AALP and CEP1 subfamilies (Fig. 2). The four *L. perenne* PLCPs R_2516 (RD21-like, subfamily I), R_4870 (AALP-like, subfamily VIII), R_182 (XCP2-like, subfamily III) and R_11872 (CTB3-like, subfamily IX) were identified as closely related to the well characterized maize apoplastic PLCPs CP1, CP2, XCP2 and CathB, respectively (van der Linde *et al.*, 2012*a*). Therefore, we renamed these *L. perenne* PLCPs to LpCP1, LpCP2, LpXCP2 and LpCathB (Fig. 2).

**Fig. 2.**
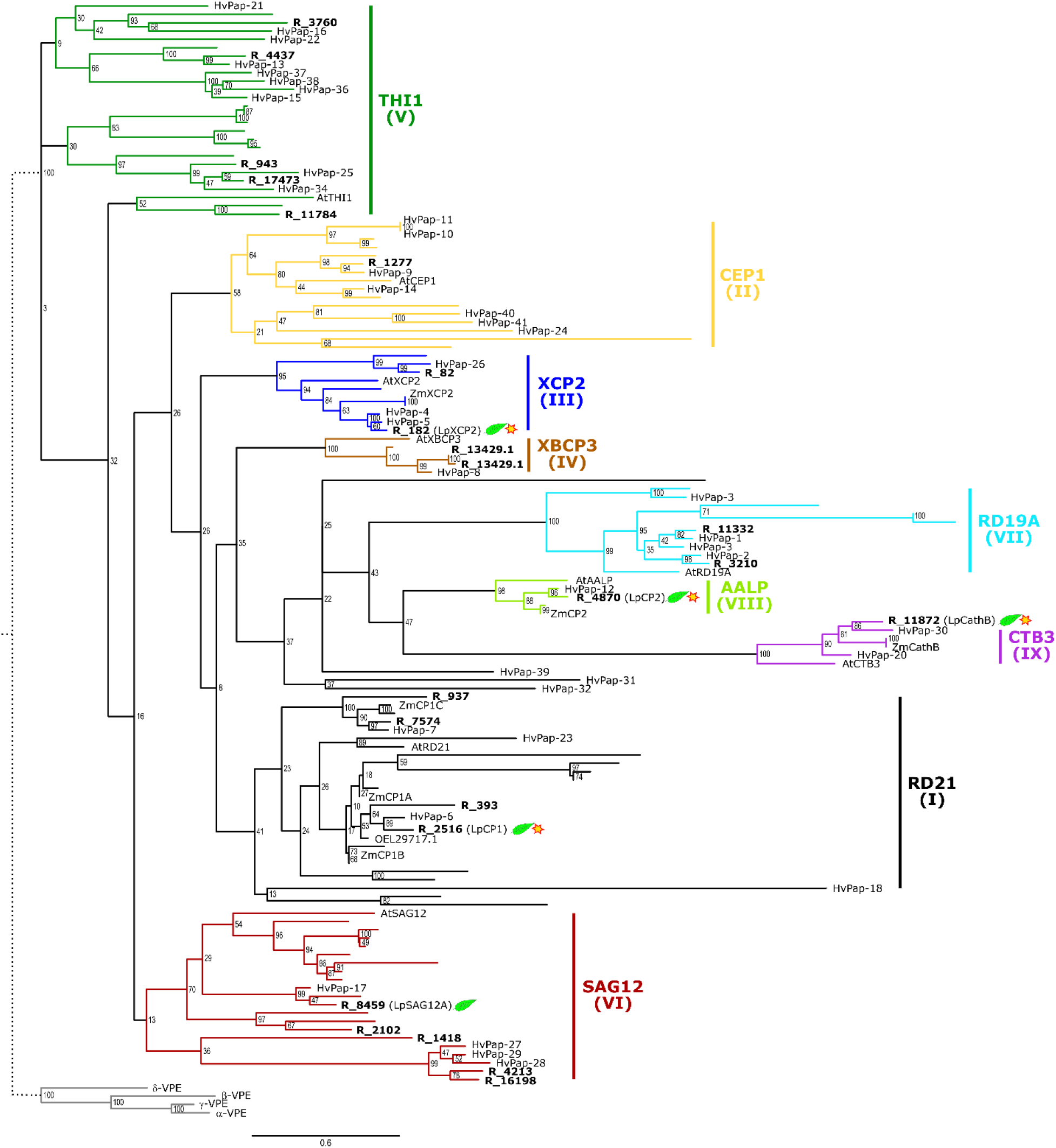
Phylogeny and subfamily classification of *L. perenne* PLCPs. Phylogenetic analysis was performed using 23 *L. perenne* PLCP sequences identified via functional domain analysis (R numbers), 52 PLCP sequences from the maize line B73 obtained from MEROPS database (www.ebi.ac.uk/merops, branches without label due to space constriction), 6 PLCP sequences from the maize line EGB, and 38 PLCP sequences of *Hordeum vulgare* (HvPaps). The four legumains (AtVPEs) from *A. thaliana* were used to root the phylogenetic three and one member of each PLCP subfamily also from *A. thaliana* (bold, colored) for the subfamily classification. In this analysis, full length sequences were used, including signal peptide, auto-inhibitory pro-domain, protease C1-domain and if present granulin domain. Leaf symbol, proteases identified in the leaf apoplast proteome analysis; star, proteases found as active enzymes in apoplast using ABPP. To fit the phylogenetic tree in the window the branch of the outgroup has been shortened (dotted line).

### Protein composition of the apoplast is altered during endophyte associations

The results of the ABPP experiment showing reduced activity of apoplastic PLCPs in *L. perenne* plants infected with the endophyte *E. festucae* and the large variation observed in the loading controls indicate that the host apoplastic proteome is significantly altered during endophyte associations. To identify changes in the proteome induced by *E. festucae* infection, apoplastic fluids of infected and uninfected plants were isolated. Quantitative apoplast proteome analysis using stable isotope labelling by reductive dimethylation identified 1153 protein groups, of which 1092 proteins originated from *L. perenne* and 86 from *E. festucae* (Suppl. Dataset S3). Of these, 572 (CT/mock) and 550 (Fl1/mock) proteins were quantified in at least 2 biological replicates for the CT/mock and Fl1/mock proteomes, respectively. Among the proteins quantified in response to CT, 552 belonged to *L. perenne* and 20 to *E. festucae* whereas in response to Fl1, 530 quantifiable proteins originated from *L. perenne* and the same 20 for *E. festucae* (Suppl. Dataset S3). Protein abundance differed depending on the endophyte inoculation. Compared to mock samples, Fl1 infected plants showed stronger changes in proteome composition than CT infected plants (Fig. 3A). These differences in proteome composition could be explained by the biomass variation in colonised tissue previously described for Fl1 in comparison to CT. Previous analyses have shown that Fl1 represents 1–2% of total biomass in infected *L. perenne* plants (Young *et al.*, 2005) whereas the biomass colonization of CT is estimated to approximately 0.2 % of leaf tissue in sheath (Tan *et al.*, 2001). Notably, 51 plant proteases were identified in the apoplastic proteome, including 7 aspartic proteases, 21 serine hydrolases, 9 cysteine proteases and 14 proteases belonging to other classes mostly metallo-peptidases (Fig. 3B and Suppl. Dataset S3). Twenty-four of these plant proteases were quantified for the Fl1/mock proteome. The cysteine protease, R_8459, a SAG12-like PLCP, was significantly reduced in the Fl1-infected proteome compared to mock. In contrast, two plant serine proteases, R_2759, an alpha/beta serine hydrolase of the peptidase S28 family and R_14255, a serine carboxypeptidase of the peptidase S10 family were significantly more abundant in response to Fl1 infection (Fig. 3C, left panel). Twenty-five plant apoplastic proteases were quantified in the CT/mock proteome although no significant reduction in their abundance was observed in response to CT infection. Notably, three proteases showed a significant accumulation: the same R_2759 and R_14255 serine proteases also found with higher abundance in response to Fl1 and R_3647, an aspartic protease with C-terminal homology to a xylanase inhibitor (Fig. 3C, right panel).

**Fig. 3.**
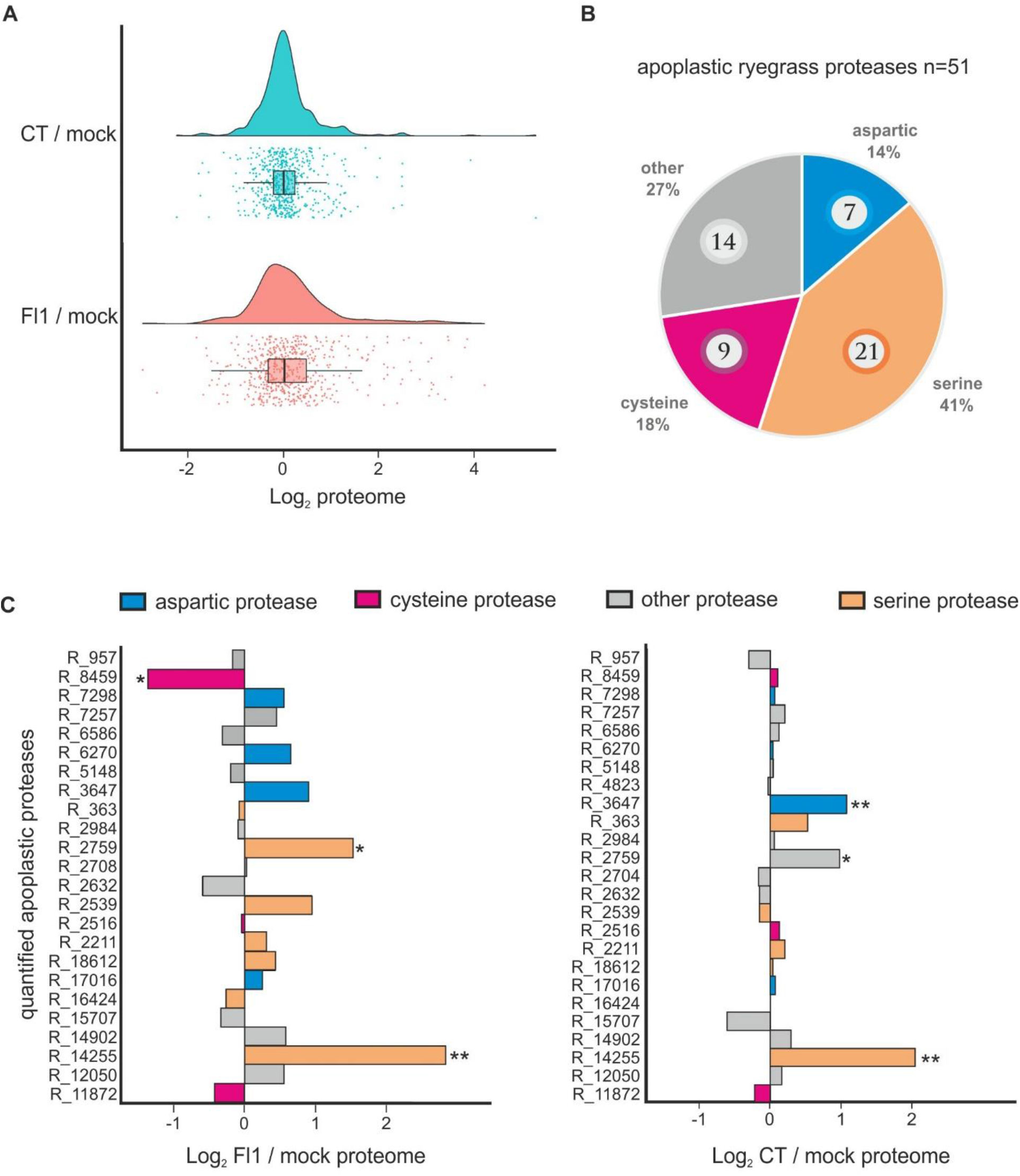
Apoplast proteome analysis of Fl1 and CT infected and mock-treated leaves. (A) Raincloud plot of Fl1/mock and CT/mock protein quantification. Apoplast proteins from Fl1, CT and mock-treated plants were stable isotope labelled by reductive dimethylation. The average log_2_ -transformed ratios Fl1/mock and CT/mock of the 530 and 552 proteins, respectively, were calculated for proteins quantified in at least two of three replicates. Average fold changes are shown as a raincloud plot combining a density graph with single dot plots for each quantified protein and a boxplot depicting the mean of the data assembly. (B) Overview of identified apoplastic proteases in the leaf proteome analysis. 51 ryegrass proteases identified in our proteome analysis were grouped according to their catalytic mechanism: aspartic proteases (blue), cysteine proteases (magenta), serine proteases (peach) and other (grey). The total number of each group of identified proteases is shown in circles. The percentage (%) of protein groups was calculated based on the total number of identified proteases. (C) Quantification of apoplastic proteases in Fl1 and CT infected leaves. Mean log_2_ -transformed ratios (n=3 biological replicates, at least quantified in 2 out of 3 replicates) are individually plotted for each of the proteases. 24 and 25 proteases were quantified for FL1/mock and CT / mock, respectively. * represent significant differences (* = p<0.05, ** = p< 0.01, LIMMA-moderated Students *t*-test).

### Identification of active PLCPs in the L. perenne apoplast

Our ABPP experiment showed that uninfected *L. perenne* plants, likely resembling *L. perenne* in its natural environment, maintain active PLCPs possibly to assist the proteolytic activity during diverse biological processes such as development, senescence, abiotic stresses and also as a response against pathogen attack. To further characterize and identify active PLCPs in the apoplast of *L. perenne* a pull-down of active PLCPs was performed using the activity-based probe DCG-04 (Greenbaum *et al.*, 2000). Apoplastic fluid of uninfected (E−) leaves was isolated and labelled with DCG-04. Biotinylated proteins were affinity purified using streptavidin beads and subjected to an on-bead digest (OBD) followed by mass spectrometry analysis. As a background control a no-probe-control proteome was equally treated with streptavidin beads. To confirm a successful affinity purification, proteins were boiled from the beads and examined by western blot analysis using streptavidin-HRP antibody. Two main signals between 25 and 35 kDa were observed representing PLCP labelling (Fig. 4A). MS analysis identified four active PLCPs present in *L. perenne* apoplast: R_11872 (LpCathB, CTB3 subfamily IX), R_2516 (LpCP1, RD21 subfamily I), R_4870 (LpCP2, AALP subfamily VIII) and R_182 (LpXCP2, XCP2 subfamily III) (Suppl. Dataset S3). LpCathB and LpCp1 are the most abundant PLCPs followed by LpCP2 and LpXCP2, estimated by their LFQ intensities (Fig. 4B; suppl. Dataset S3). All four active PLCPs found in the apoplast of *L. perenne* belong to different subfamilies and share the same domain structures containing a signal peptide, pro-domain and mature protease with the catalytic triad Cys, His and Asn. LpCP1 additionally contains a proline-rich repeat and a granulin domain (Fig. 4B). Interestingly, the PLCP SAG12 (R_8459) which was found to be significantly less abundant in the Fl1 sample (Fig. 3C) was not identified as an active PLCP in the mock sample suggesting that the loss of activity observed for the Fl1 treated samples in the ABPP experiment correspond to the active PLCPs LpCP1, LpXCP2, LpCP2 and LpCathB.

**Fig. 4.**
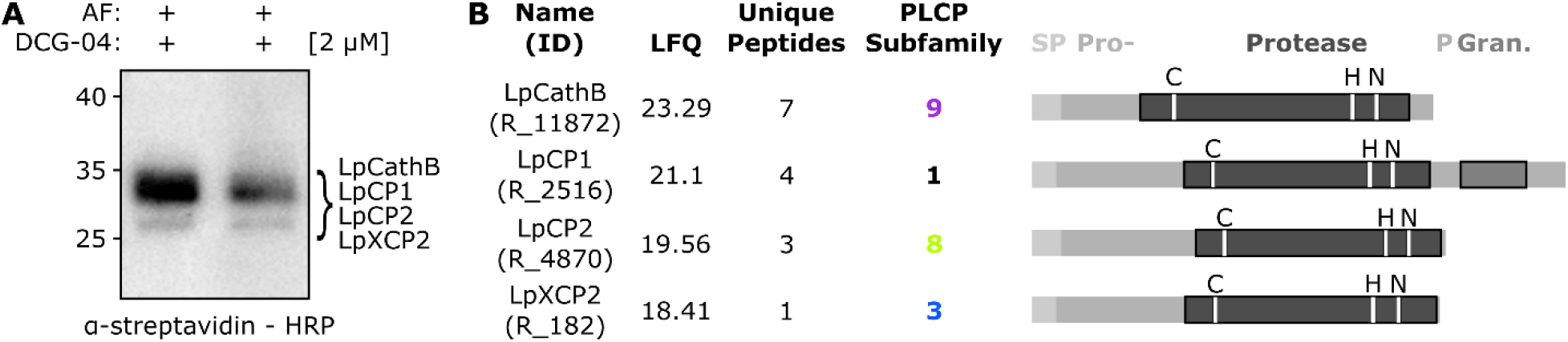
Identification of active *L. perenne* apoplastic PLCPs. (A) Pull-down of apoplastic PLCPs using DCG-04. Apoplastic fluid (AF) of endophyte free plants was labelled with the biotinylated probe DCG-04. A background control without DCG-04 (no-probe control, NPC) was performed. Labelled proteins were purified using streptavidin beads. Biotinylated proteins were detected using α-streptavidin-HRP antibody. Shown is a representation of two biological replicates. Estimated sizes for *L. perenne* (Lp) PLCPs: LpCathB, LpCP1, LpCP2 and LpXCP2 are shown in brackets. (B) Schematic representation of identified apoplastic PLCPs of *L. perenne*. Pull – down samples were subjected to on bead digest (OBD) and subsequent mass spectrometry analysis. Samples were analysed in triplicates and only proteases present in at least two of the three replicates were considered. SP = signal peptide, Pro- = auto-inhibitory prodomain, Protease = protease C1-domain, P = Proline-rich domain, Gran = granulin domain. Letters above the protease domain represent the catalytic triad (C, cysteine; H, histidine; N, asparagine). Coloured numbers show the PLCP subfamily. LFQ, MaxQuant-label-free quantification intensity as a measure of abundance.

### A cysteine protease inhibitor is present in the apoplast of Fl1 infected leaves

Since apoplastic PLCP activity but not the abundance of the majority of cysteine proteases was strongly reduced in response to *E. festucae* infection (Fig.1, 3C), we hypothesized that inhibitor molecules modulate PLCPs during endophytic *E. festucae* colonization. To test this assumption, a convolution ABPP was performed. Two apoplastic fluids (E− and E+) were combined prior to labelling with the activity-based probe (BL). If one of the apoplastic fluids (E+) contains a PLCP inhibitor in excess, the inhibitor will suppress active PLCPs present in the other apoplastic fluid (E−). To ensure that a reduction in PLCP activity is not caused by dilution, the two apoplastic fluids were also mixed after labelling (AL). Thus, monitored PLCP activity should be an average of the individual PLCP activities, when combined after labelling. A reduction of protease activity in BL compared to AL would indicate the presence of excess inhibitor in one of the apoplastic fluids (Fig. 5A;Chandrasekar *et al.*, 2017). The convolution experiment was performed with MV202 labelled E− and E+ apoplastic fluids. E-64 pre-incubated samples served as control to ensure that observed signals were specific to the MV202 labelling and corresponded to PLCPs. The convolution ABPP revealed that combining mock (E−) and infected (E+) apoplastic fluid before labelling (BL) caused a stronger reduction in PLCP activity compared to the combination after labelling (AL; Fig. 5B). Notably, a strong signal at ca. 35 kDa was observed in the SyproRuby gel for the E+ sample. We hypothesized that this signal could represent the stabilization of PLCPs by an inhibitor. Fluorescent signal quantification of three biological experiments was performed and confirmed that the PLCP activity of BL was halved compared to AL (Fig. 5C), indicating that reduction in PLCP activity in response to *E. festucae* interaction is caused by an apoplastic PLCP inhibitor.

**Fig. 5.**
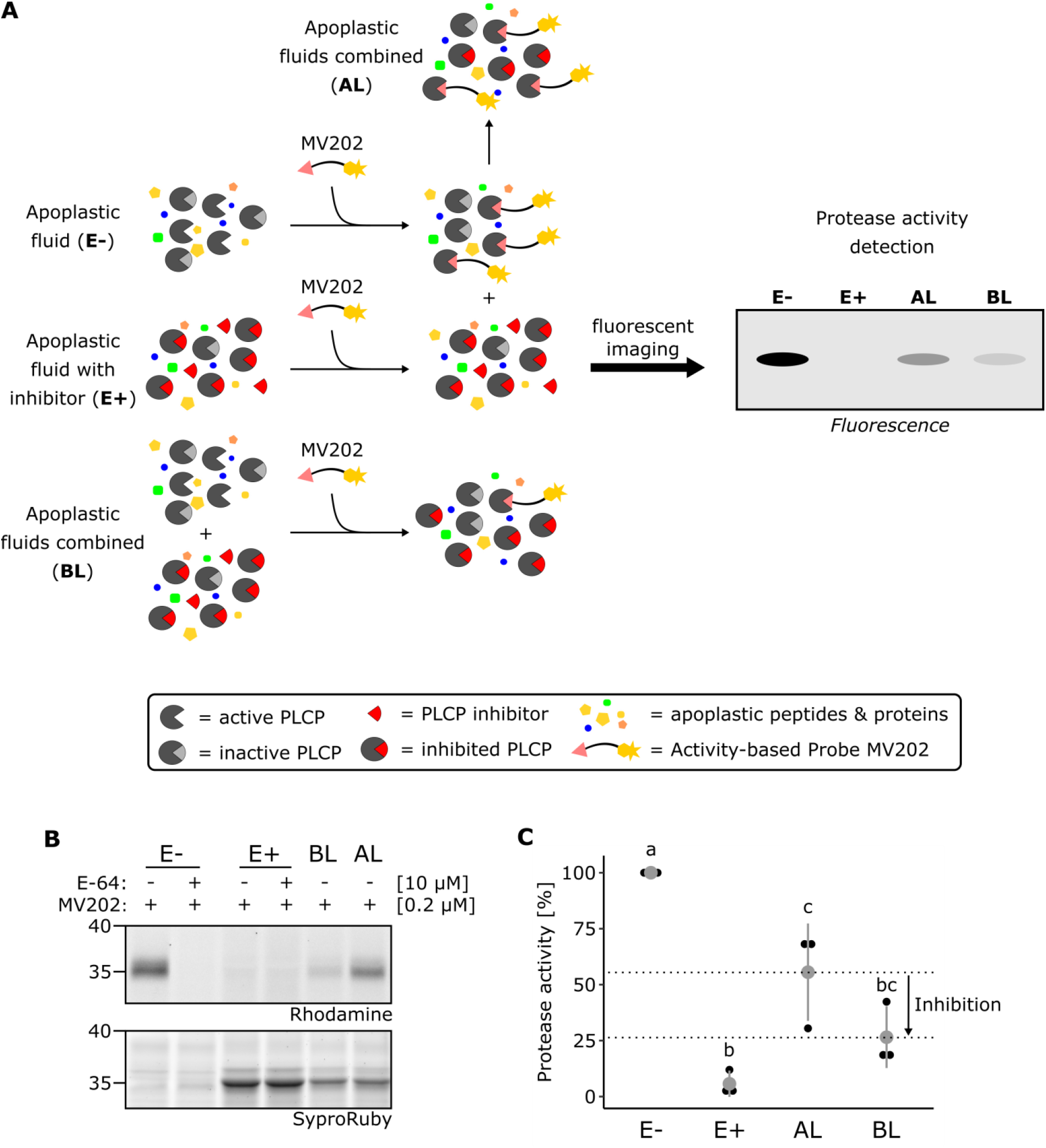
An apoplastic PLCP inhibitor is present in *E. festucae* infected plants. (A) Convolution ABPP workflow. Apoplastic fluid with (E+) and without (E−) potential PLCP inhibitor is mixed in a 1:1 ratio, allowing the potential PLCP inhibitor in E+ to inhibit active PLCPs in E− before labelling (BL). All three fluids (E−, E+, BL), were labelled with the activity-based probe MV202. As control, E+ and E− are mixed (1:1) after labelling (AL). The fluorescent signals detected in AL should represent the average signals of E+ and E−. If excess inhibitor is present in E+ apoplastic fluid, signal intensities of BL will be lower than signal intensities of AL. Figure was adapted from Chandrasekar *et al.*, 2017. (B) An apoplastic PLCP inhibitor is produced during *E. festucae* infection. Apoplastic fluid of E+ and E-plants was isolated and pre-incubated with E-64 or the equivalent amount of DMSO followed by labelling with MV202. One volume of E+ fluid and one volume of E− fluid were incubated for 45 min prior to the labelling with MV202 (BL). After labelling E+ and E− were mixed in a 1:1 ratio (AL). Samples were separated via SDS-PAGE and labelled PLCPs were visualized by in-gel fluorescent scanning using a rhodamine filter. Sample loading was visualized via SyproRuby staining. (C) Quantification of convolution ABPP. Labelling intensities were quantified and normalized to samples treated with E-64. AL and BL were normalized to the average of E-64 treated samples of E+ and E−. Protease activity of E− was set to 100% and PLCP activity was calculated in relation to the E-sample. The red dot represents the mean of three independent biological replicates (black dots), while the red line represents the standard deviation. Different letters indicate significant differences between the means according to Tukey’s test (*P* < 0.05).

### Mining the apoplastic proteome for potential PLCP inhibitors

To investigate if the potential apoplastic PLCP inhibitor is of endophyte origin, we screened the *E. festucae* Fl1 genome for orthologues of the known plant pathogen - derived PLCP inhibitors Avr2 (*Passalora fulva*), EpiC1 and Avrblb2 (*Phytophthora infestans*), VAP1 (*Globodera rostochiensis*), Pit2 (*Ustilago maydis*), popP2 (*Ralstonia solanacearum*), SDE1 (*Candidatus Liberibacter asiaticus*) and Cip1 (*Pseudomonas syringae* pv. tomato DC3000) (Suppl. Table S4). A blastp analysis revealed two potential VAP1 orthologs, Fl1_004109 and Fl1_007387 with 20.1% and 12.04% identity to GrVAP1, respectively. These two candidates were the only hits displaying E-values with significant homology (Suppl. Table S4). In a second, unbiased, approach our generated apoplastic proteome dataset was screened for the presence of Fl1 *E. festucae* proteins. This approach identified 86 Fl1 proteins of which 22 did not show homology to a known PFAM domain and represent proteins with unknown function. Of those 86, twenty proteins could be quantified since they were present in at least two biological replicates. A functional annotation using InterProScan (www.ebi.ac.uk/interpro), prediction tools for secretion signal (apoplastP 1.0, http://apoplastp.csiro.au) and a prediction of a virulence function (EffectorP 1.0, http://effectorp.csiro.au) was performed (Suppl. Dataset S3). From those Fl1 quantified proteins only one, Fl1_003471, was predicted as an effector with apoplastic localization which is more abundant in CT samples than in Fl1 samples (Suppl. Table S5 and suppl. Dataset S3). From the 66 non-quantified proteins, 17 were of unknown function and therefore functionally annotated as previously described (Suppl. Table S5). Fl1_002869, Fl1_003333 and Fl1_005240 were predicted as putative apoplastic effectors. None of the identified apoplastic *E. festucae* Fl1 proteins showed homology to any cysteine protease inhibitor and the two orthologues of VAP1 were not found in our generated proteome dataset.

Since our analysis of the apoplastic proteome did not identify *E. festucae* effector candidates with significant similarity to known PLCP-inhibitors, we screened the apoplastic proteome dataset for potential PLCP inhibitors from the host plant. Seven putative cysteine and serine protease inhibitors of plant origin were identified: three cystatins R_12141, R_2071 and R_27228 and four serine-type peptidase inhibitors (Table 1). Of these, one cystatin (R_12141) was accurately quantified in two of the three experiments and did not show a significant difference in abundance between endophyte (Fl1 or CT) and mock inoculated samples (Suppl. Dataset S3). A phylogenetic analysis using plant cystatins from *A. thaliana* (Arabidopsis), *H. vulgare* (barley), *Z. mays* (maize) and *O. sativa* (rice) as the chicken (*Gallus gallus*) egg white cystatin (CEWC) showed three main clusters: cluster I represents only monocot cystatins, cluster II with four Arabidopsis members and representative cystatins of different monocots and cluster III with representative sequence homologues of AtCYS-2. R_12141 and R_2071 belonged to cluster III whereas R_27228 grouped to the monocot cluster I (Fig. 6). The closest homologue of R_12141 is HvCPI2, which is involved in barley defence against *Magnaporthe oryzae* (Velasco-Arroyo *et al.*, 2018). R_12141 also clustered with the *A. thaliana* cystatin AtCys-1 and the maize cystatin CC1, and therefore this protein sequence was named LpCys1. For R_2071 the closest homologue is the barley cystatin HvCPI4 and AtCys-6. Considering the close phylogeny proximity of R_27228 to the maize cystatin CC9 we have named this sequence LpCys9 (Fig. 6). In summary, in our proteome dataset we could not identify apoplastic proteins from *E. festucae* with annotated function as protease inhibitors. Nevertheless, we have identified three plant cystatins present in the apoplast of infected samples which could be manipulated by *E. festucae* to inhibit plant PLCPs thus promoting infection and a successful colonization.

**Table 1.** Identified apoplastic inhibitors from plant origin. Shotgun MS analysis of Fl1, CT and mock isotopically labeled apoplastic proteomes screened for the presence of protease inhibitors.

| ID | Inhibitor annotation | unique peptide | Sequence coverage [%] | MQ score |
| --- | --- | --- | --- | --- |
| R_12141 | Cystatin domain (IPR000010) * | 2 | 12.2 | 22.9 |
| R_196 | Proteinase inhibitor I13 ** | 2 | 54.3 | 3.6 |
| R_2071 | Cystatin domain (IPR000010) * | 2 | 8.9 | 10.1 |
| R_2240 | Proteinase inhibitor I13 ** | 1 | 28.2 | 188.6 |
| R_27228 | Cystatin domain (IPR000010) * | 2 | 15.2 | 4.9 |
| R_28259 | Proteinase inhibitor I12, Bowman-Birk type ** | 2 | 17.1 | 51.9 |
| R_386 | Bowman-Birk type proteinase inhibitor (IPR035995) ** | 1 | 10.0 | 25.0 |

Inhibitor identity (ID).
Inhibitor annotation based on the uniprot database (www.uniprot.org).
MaxQuant score (MQ score).
\* Cysteine protease inhibitor family (cystatin domain).
\*\* Serine proteinase inhibitor family

**Fig. 6.**
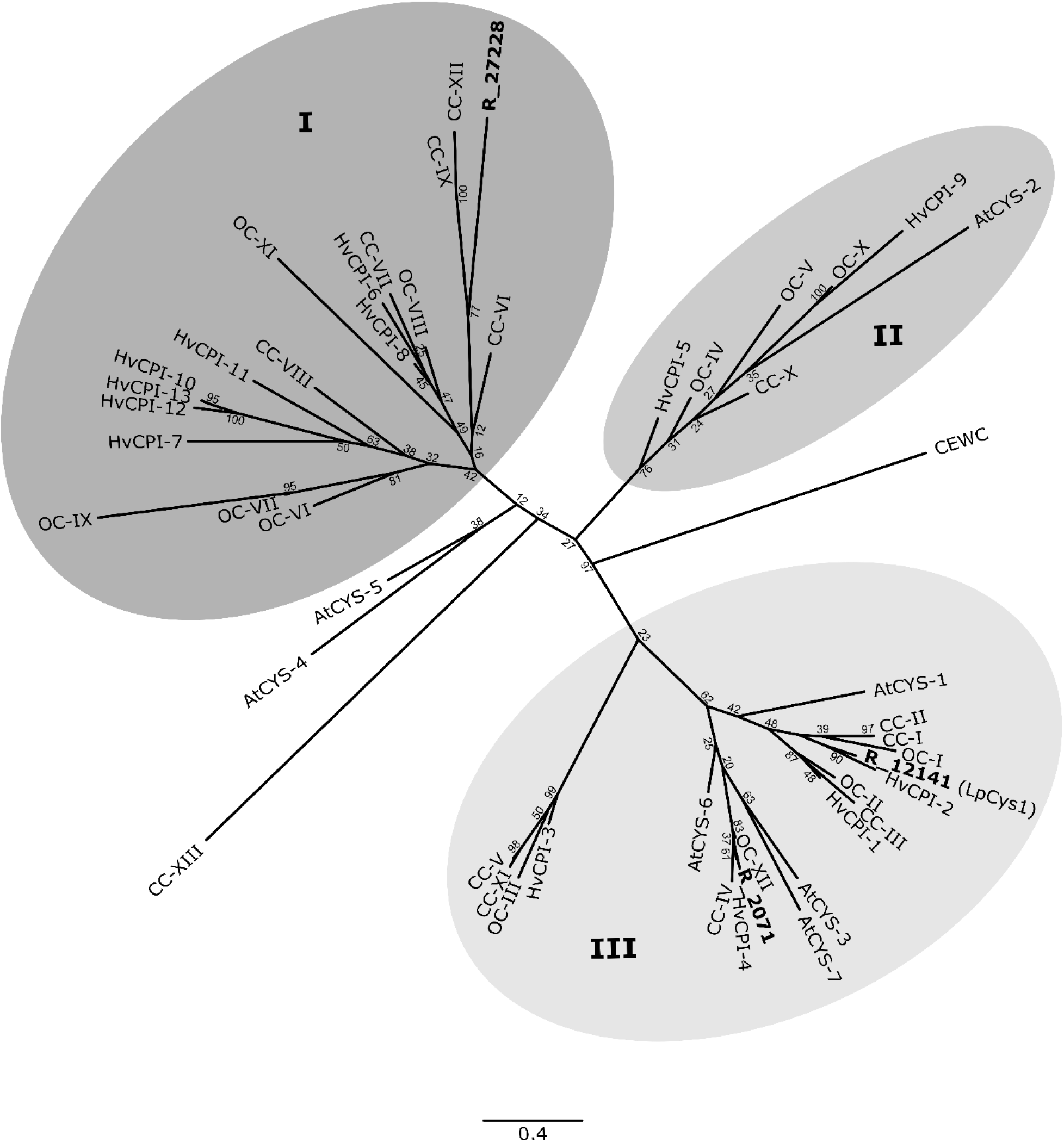
Phylogentic analysis of identified cystatins from *L. perenne.* Alignment of full-length sequences of *L. perenne* R_12141 (LpCys1), R_2071 and R_27228 (LpCys9) as well as *H. vulgare* (HvCPI), *Z. mays* (CC), *O. sativa* (OC), *A. thaliana* (AtCYS) and the chicken cystatin (CEWC) were generated using MAFFT. The unrooted radial tree was generated with RAxML (v8.2.12). 100 bootstraps were performed and bootstrap values are indicated. The tree was visualized with FigTree. Grey circles show the main three clusters (I, II and III) obtained for the tested cystatins.

### The L. perenne cystatin LpCys1 inhibits LpCP2

To further study the role of *L. perenne* cystatins in the inhibition of plant PLCPs we selected LpCys1 since it was the only reliably quantified cystatin in our apoplast proteome. LpCys1 is closely related to the barley cystatin HvCPI2, which has been shown to strongly inhibit the barley PLCPs HvPap-6 and HvPap-10 (Martinez *et al.*, 2009). GST-tagged LpCys1 was expressed in *E. coli* and purified by affinity chromatography followed by gel filtration. Purified LpCys1 was tested for its inhibitory capacity on the four apoplastic *L. perenne* PLCPs: LpCP1, LpCP2, LpXCP2 and LpCathB which were heterologous expressed in *N. benthamiana* leaves using Agrobacterium transient transformation. PLCP activity was determined from *N. benthamiana* apoplastic fluids containing the PLCPs using the activity based probe MV201 (Richau *et al.*, 2012). A concentration range (0 to 4.5 μM) of purified LpCys1 was used for inhibition assays with the four PLCPs. Commercially available chicken cystatin (CEWC) was used as a positive control. Three controls were used in this ABPP experiment: E-64 to test for the specificity of MV201 signals, *N. benthamiana* expressed CP1Amut (a catalytic inactive maize PLCP, Schulze Hüynck *et al.*, 2019), as negative control for the PLCP background in *N. benthamiana* apoplastic fluids and a no-probe control (NPC), to detect unspecific fluorescent background. The activity of LpCP1 increased with low concentrations of LpCys1, reaching a maximal peak at ca. 2.5 μM of incubation with the cystatin. With cystatin concentrations greater than 2.5 μM the activity of LpCP1 decreases in a concentration dependent manner (Fig. 7A, B). SyproRuby staining showed a band at ca. 26 kDa, likely representing LpCP1, which increases in intensity until the addition of ca. 2.5 μM LpCys1 but continues stable with increasing concentrations of the cystatin (Fig. 7A). Notably, signal quantification from four biological replicates confirmed a six-fold increase of LpCP1 activity after incubation with 2.5 μM LpCys1 and further increasingly concentrations of the cystatin decreases LpCP1 activity (Fig. 7B). In contrast, CEWC showed a strong inhibition against LpCP1 already at 0.5 μM (Fig. 7B and suppl. Fig. S2). In case of LpCP2, LpCys1 showed a concentration dependent inhibitory effect, resulting in complete inactivation of the protease already at 3 μM LpCys1 (Fig. 7C). Inhibition of LpCP2 was also observed by CEWC, although with a much weaker inhibitory capacity than LpCys1 (Fig. 7D & Suppl. Fig. S2). LpXCP2 was poorly inhibited by both, LpCys1 and CC, and ca. 50% of the inhibition was reached with 4.5 μM, the maximum tested concentration for both inhibitors, suggesting that LpCys1 has a poor affinity towards LpXCP2 (Fig. 7E-F and Suppl. Fig. S2). Finally, LpCys1 did not inhibit LpCathB although CEWC showed a concentration dependent inhibitory effect suggesting that LpCathB might not be a target for LpCys1 (Fig. 7G-H & Suppl. Fig. S2). These results suggest that LpCys1 has a stronger inhibitory capacity against LpCP2 than towards other PLCPs. Thus, LpCys1 is a potent and specific inhibitor of LpCP2 that might contribute to the general reduction of apoplastic PLCP activity during *E. festucae* colonization. However, since only one out of four active proteases is sensitive to this cystatin, it is likely that additional inhibitors are involved in PLCP inactivation during this fungal interaction.

**Fig. 7.**
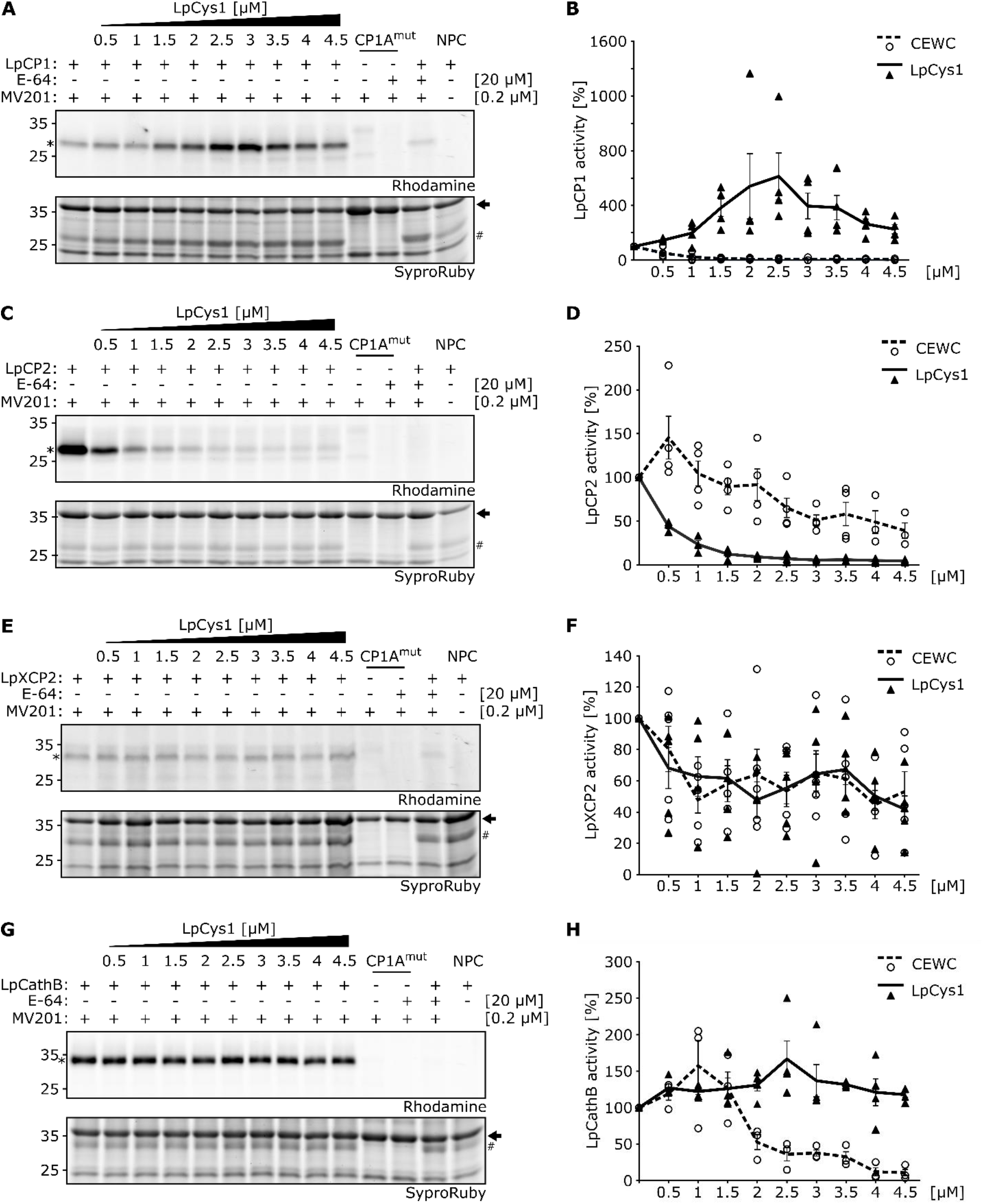
Inhibitory activity of LpCys1 on overexpressed ryegrass PLCPs. Four PLCPs of *L. perenne* were overexpressed in *N. benthamiana* using Agrobacterium-mediated transformation. Apoplastic fluid containing PLCPs was isolated and monitored using the fluorescent probe MV201. Samples were pre-incubated for 15 min with 20 μM E-64 or a concentration range (0 – 4.5 μM) of heterologous expressed LpCys1 or CWEC followed by 2 h labelling with 0.2 μM MV201. As background controls, samples containing CP1A^mut^, an inactive maize PLCP, and the no-probe control (NPC) were prepared. The activity and inhibitory effect of LpCys1 on LpCP1 (A), LpCP2 (C), LpXCP2 (E) and LpCathB (G) was analysed using in gel fluorescent scanning and as a loading control SyproRuby staining was performed. For each analysed PLCP (marked as asterisk) signals were quantified and normalized to a loading control signal (marked with an arrow) and activity without inhibitor was set to 100%. Normalized activity values [%] for LpCP1 (B), LpCP2 (D), LpXCP2 (F) and LpCathB (H) were plotted against the concentration [μM] of LpCys1 (solid line) and CWEC (dotted line). Error bars represent the standard error of three biological replicates.

## Discussion

In this study we have shown that apoplastic papain-like cysteine proteases are inhibited during endophytic colonisation of the *L. perenne* ryegrass, similar to pathogen colonisation. We identified the PLCPs LpCP1, LpCP2, LpXCP2 and LpCathB being active in the apoplast of uninfected *L. perenne* plants. Interestingly, these PLCPs classify into different PLCP subfamilies, known to be hubs in plant immunity (Misas Villamil *et al.*, 2016). The most abundant of the four active apoplastic PLCPs in uninfected *L. perenne* leaves was LpCathB. In both, *N. benthamiana* and *A. thaliana* homologues of LpCathB are involved in the hypersensitive response triggered by bacterial avirulent pathogens (Gilroy *et al.*, 2007; McLellan *et al.*, 2009; Ge *et al.*, 2016) although it might not function as a universal regulator of the hypersensitive response (Thomas and van der Hoorn, 2018). The second most abundant active apoplastic PLCP was LpCP1, followed by LpCP2 and LpXCP2. All three proteases have orthologues in maize which were found to be activated in the apoplast after salicylic acid treatment and are involved in the defence response against the biotrophic fungus *U. maydis* (van der Linde *et al.*, 2012*a*; Mueller *et al.*, 2013). Maize CP1 and CP2 apoplastic proteases are targeted and inhibited by the *U. maydis* effector Pit2 which acts as a substrate mimic molecule to achieve a successful inhibition (Mueller *et al.*, 2013; Misas Villamil *et al.*, 2019). LpCP1 belongs to the RD21-like subfamily (I) and contains a granulin domain, which is exclusively found in members of subfamily I or IV (Richau *et al.*, 2012; Misas Villamil *et al.*, 2016). Members of this subfamily have been described to play a crucial role during pathogen attack. The tomato C14 protease is inhibited during *Phytophthora infestans* infection by the cystatin-like effector proteins EpiC1 and EpiC2B (Kaschani *et al.*, 2010) and by the chagasin-like Cip1 inhibitor during Pseudomonas infection (Shindo *et al.*, 2016). Additionally, the RxLR effector of *P. infestans* targets C14 to prevent its secretion into the apoplast (Bozkurt *et al.*, 2011). Moreover, in barley, which is phylogenetically closely related to ryegrass, HvPap-6 accumulated after *Magnaporthe oryzae* treatment, particularly at late stages of infection and after infestation with the mite *Tetranychus urticae* where mostly the pre-mature form of HvPap-6 accumulated (Diaz-Mendoza *et al.*, 2017). LpCP2 belongs to the AALP-like subfamily (VIII) and like its closest homologue in barley, the thiol protease aleurain Hv-Pap12, LpCP2 is also present and active in barley leaf extracts (Frank *et al.*, 2019). Similar to LpCP1 subfamily members, LpCP2 orthologues play a role in defence against different pathogens. Silencing of CYP1/2 in *N benthamiana* increases susceptibility against *Colletotrichum destructivum* (Hao *et al.*, 2006). In maize, CP1 and CP2 have been shown to function in the release of Zip1, a peptide signalling molecule that activates salicylic acid immune responses (Ziemann *et al.*, 2018). Together, this body of evidence indicate that members related to these PLCP subfamilies need to be shut-down by pathogens to avoid activation of plant immunity.

Do endophytes need to apply similar strategies on modulation of PLCPs as pathogens? In *E. festucae* Fl1 and CT infected *L. perenne* plants the abundance of PLCPs did not significantly differ from mock plants, although PLCP activity was almost fully diminished, indicating that the reduction in PLCP activity is caused via inhibition rather than protein degradation. Based on the results presented here, we speculate two ways of inhibition: the putative inhibitor could be of plant origin, manipulated by *E. festucae* to achieve a successful colonization, or of fungal origin, a secreted effector molecule. An example of an inhibitor of plant origin is the cystatin CC9 which was identified as an important compatibility factor in the maize – *U. maydis* interaction. Upon *U. maydis* infection *cc9* expression is induced and CC9 suppresses plant immunity via PLCP inhibition, thus enabling *U. maydis* colonization (van der Linde *et al.*, 2012*a*). CC9 inhibits all apoplastic PLCPs and is required for early stages of *U. maydis* colonization since at later stages of infection the Pit2 inhibitor likely takes over as a more specific inhibitor of the apoplastic proteases CP1 and CP2 (van der Linde *et al.*, 2012*a*; Misas Villamil *et al.*, 2019). Interestingly, the mechanism of inhibition of CP1 and CP2 by *U. maydis* Pit2 resembles the “activation – inhibition” of LpCys1 towards LpCP1 and LpCP2. Whilst in maize CP2 is inhibited by Pit2, CP1 is first stabilized leading to an increased CP1 activity and eventually, at higher concentrations of Pit2, to inhibition (Misas Villamil *et al.*, 2019). In case of the *L. perenne – E. festucae* interaction, LpCys1 did not efficiently inhibit LpCP1 but rather activates it, contrary to LpCP2 where LpCys1 achieves a full inhibition. These results suggest that LpCys1 is likely stabilizing LpCP1 until the batch of available zymogen has been consumed and the inhibition can then take place in a concentration dependent manner.

LpXCP2 and LpCathB are not inhibited by the cystatin LpCys1. These results indicate that another inhibitor besides LpCys1 is involved in the PLCP inhibition in response to *E. festucae* interaction. CC9 is not the closest orthologue of LpCys1 but of LpCys9 which might therefore represent an interesting candidate for ryegrass PLCP inhibition. If LpCys9 expression is similarly to CC9, one could speculate that it might be transiently activated at early stages of infection. In this study we did not examine early stages of infection which might be reflected in the innermost leaf blade of the pseudostem tissue (Schmid *et al.*, 2016), nevertheless a strong PLCP inhibition was observed during endophytic interactions indicating the production of a PLCP inhibitor also in a long term systemic host colonization. These findings confirm that endophyte infections of *L. perenne* lead to major alterations of the host metabolism, development and apoplastic proteome (Scott *et al.*, 2018; Green *et al.*, 2020).

The presence of LpCys1 in the apoplast could have an alternative function unrelated to the inhibition of PLCPs. In barley, the HvCPI-2 cystatin, the closest orthologue to LpCys1, showed a strong fungicide effect against *Botrytis cinerea* and *Fusarium oxysporum* mycelia (Abraham *et al.*, 2006), suggesting that the presence of LpCys1 could be part of the plant immune response against *E. festucae*, rather than LpCys1 being manipulated by *E. festucae* to facilitate PLCP inhibition. Indeed, one of the most highly expressed fungal genes *in planta* is a chitinase (Eaton *et al.*, 2010), suggesting host defence responses are activated and *E. festucae* evades plant immunity by altering or masking the chitin composition of hyphae *in planta* (Becker *et al.*, 2016).

Based on our results it is possible that the inhibition of ryegrass PLCPs during mutualistic interactions is the result of a cooperative effect of plant cystatins and a secreted effector from *E. festucae*. We have identified four putative effector candidates present in our apoplast analysis and two VAP-1 orthologues that could potentially inhibit *L. perenne* PLCPs. Both *E. festucae* VAP1-like proteins match to an allergen V5/SCP domain containing protein of *Claviceps purpurea* and *Moelleriella libera* based on the uniprot database (www.uniprot.org). Fl1_004109 also has a ‘hit’ with a basic form of pathogenesis protein 1 of *Pochonia clamydospora*. Notably, all best ‘hits’ correspond to proteins restricted to the order Hypocreales (*Claviceps spp*., *Metarhizium spp*., *Pochonia spp*., *Moelleriella spp*., *Ustilaginoiea spp*.) suggesting that both *E. festucae* identified VAP1 orthologues might be ubiquitous proteins of the family Clavicipitaceae and not specific effectors from *E. festucae*. Notably, common features can be found for cystatins such as the Q-V-G motif (Gln-Xaa-Val-Xaa-Gly), a Pro -Trp or Leu -Trp dipeptide motif in the C-terminal region and a conserved Gly residue in the N-terminal region (Benchabane *et al.*, 2010). These common features are not found in pathogen – derived PLCP inhibitors suggesting that pathogens evolve independent strategies to suppress protease activity making challenging bioinformatic searches of this type of effector molecules. Whether the new potential *E. festucae* effector candidates contribute to the full inhibition during Fl1 interactions remains to be elucidated.

In summary, we have shown that during the *L. perenne* – *E. festucae* interaction, microbial endophytes modulate essential components of the plant immune system, similar to pathogenic interactions. In this case, the inhibition of apoplastic cysteine proteases might be essential and required for *E. festucae* to maintain a mutualistic interaction with its host.

## Supplementary data

Table S1: oligonucleotides

Table S2: strains

Table S3: original identifiers of *L. perenne*

Table S4: Functional annotation of “unknown” Fl1 apoplastic proteins

Table S5: screen of PLCP-inhibitor orthologs in *E. festucae* Fl1 strain

Fig. S1: Phylogenetic tree of *L. perenne* PLCPs

Fig. S2: Concentration range of the inhibition of *L. perenne* PLCPs by CEWC

Dataset S1: sequences used for PLCP phylogenetic analysis

Dataset S2: sequences used for cystatin phylogenetic analysis Dataset S3: apoplastic proteomics, annotation and functional analysis

## Acknowledgements

We would like to thank Yvonne Becker for providing the infected plants, for reading the manuscript and for giving us helpful comments and suggestions. We also thank Hermen Overkleeft (Leiden University) for kindly providing us with the ABPs. Many thanks to Ute Meyer for fruitful discussions and technical support. Also, thanks to David Winter (Massey University) for providing the *E. festucae* Fl1 protein dataset. Work in the Scott laboratory was supported by a grant (RM19009) from the Tertiary Education Commission to the Bio-Protection Research Centre, B.S. was supported by an Alexander von Humboldt Research Award.

## Author contributions

A.P. and J.C.M.V. wrote the manuscript with input from all authors. G.D., J.C.M.V., B.S., P.F.H and A.P. designed the experiments. A.P., K.G, and J.C.M.V. performed the biochemical characterization of ryegrass PLCPs. J.R.L.D and A.P. made the bioinformatic and phylogenetic analyses of plant PLCPs and cystatins. F.D. and P.F.H. performed and analysed the mass spectrometry experiments.

## Data availability statement

All data supporting the findings of this study are available within the paper and within its supplementary materials published online. All MS-based proteomics data have been deposited to the ProteomeXchange Consortium via the PRIDE (Perez-Riverol *et al.*, 2019) partner repository with the identifiers PXD022007 (reviewer login, password: ECUnxNOH) for the ABPP dataset and PXD022009 (reviewer login, password: zQhfJ1pi) for the apoplast proteome dataset.

## Notes

### Competing Interest Statement

The authors have declared no competing interest.

