## Supplementary tables & figures for "*Epichloë festucae* in mutualistic association with *Lolium perenne* suppresses host apoplastic cysteine protease activity"

**Supplementary Table S1:** oligonucleotides used in study.

| ID: | description | Sequence (5‘-3‘): | Used for: |
|---|---|---|---|
| **5104** | LpCP1_Fw1 | TTTGGTCTCAACATAATGAGGACCTCCACGGCTCTCCTG | Modular Cloning (domestication) |
| **5105** | LpCP1_Rv1 | TTTGGTCTCAACAAGAACACCGCGAACCGGAGC | Modular Cloning (domestication) |
| **5106** | LpCP1_Fw2 | TTTGGTCTCAACATGTTCATGGACAACCTCCGCTACG | Modular Cloning (domestication) |
| **5107** | LpCP1_Rv2 | TTTGGTCTCAACAAAAGCGTTCGCGCCAGTCTTTAACGGG | Modular Cloning (domestication) |
| **5108** | LpCP1_Fw3 | TTGAAGACAAAATGAGGACCTCCACGGCTCTCCTG | Modular Cloning (pICH41308) |
| **5941** | LpCP1_Rv3 | TTGAAGACAAAAGCTCAGTTCGCGCCAGTCTTTAACG | Modular Cloning (pICH41308) |
| **5094** | LpCP2_Fw1 | TTTGGTCTCAACATAATGGCCCACGGCCGCATC | Modular Cloning (domestication) |
| **5095** | LpCP2_Rv1 | TTTGGTCTCACGTCTCCGGGAGCGCGCTGGC | Modular Cloning (domestication) |
| **5096** | LpCP2_Fw2 | TTTGGTCTCAGACGAAAGACTGGAGGGAGACTGGGATCGTTAG | Modular Cloning (domestication) |
| **5097** | LpCP2_Rv2 | TTTGGTCTCAACAAGTAGACCTCCGTTGCATCCAAAATTGTTGTAT | Modular Cloning (domestication) |
| **5098** | LpCP2_Fw3 | TTTGGTCTCAACATCTACCATCCCAGGCATTTGAGTACATC | Modular Cloning (domestication) |
| **5099** | LpCP2_Rv3 | TTTGGTCTCAATCTTCACCCCAGTCGGCTCCCCAT | Modular Cloning (domestication) |
| **5100** | LpCP2_Fw4 | TTTGGTCTCAAGATGGTTACTTCAAGATGGAGCTGGGAAA | Modular Cloning (domestication) |
| **5101** | LpCP2_Rv4 | TTTGGTCTCAACAAAAGCTGCCGCCAAGATAGGGTAGG | Modular Cloning (domestication) |
| **5102** | LpCP2_Fw5 | TTGAAGACAAAATGGCCCACGGCCGCATC | Modular Cloning (pICH41308) |
| **5942** | LpCP2_Rv5 | TTGAAGACAAAAGCTCATGCCGCCAAGATAGGGTAGG | Modular Cloning (pICH41308) |
| **5116** | LpXCP2_Fw | TTGAAGACAAAATGGCTCACCCAATGAAGCTCTC | Modular Cloning (pICH41308) |
| **5945** | LpXCP2_Rv | TTGAAGACAAAAGCTCAGTTGTCCTTGGTCGGGTAGGAGGC | Modular Cloning (pICH41308) |
| **5118** | LpCath_Fw | TTGAAGACAAAATGAGGGGCGCCTCGCTTCTG | Modular Cloning (pICH41308) |
| **5940** | LpCath_Rv | TTGAAGACAAAAGCTCAGAGTATAGCAGTTCCAAAACC | Modular Cloning (pICH41308) |
| **5787** | LpCys1_Fw | TCCGAGCTCGAGATCTGCAGAATGGAGGTTTGGAAATATCG | cloning to pRSET with PvuII |
| **5788** | LpCys1_Rv | TTCGAATTCCATGGTACCAGGGCGCTTGCACCGCCCTC | cloning to pRSET with PvuII |

**Supplementary Table S2:** Strains used in this study.

| **Plasmid:** | **Background:** | **Resistance:** | **Used for:** |
|---|---|---|---|
| pL1M-F1-LpCP1::2x35S | GV3101 | Rif, Gent, Carb | for *N. benthamiana* expression |
| pL1M-F1-LpCP2::2x35S | GV3101 | Rif, Gent, Carb | for *N. benthamiana* expression |
| pL1M-F1-LpXCP2::2x35S | GV3101 | Rif, Gent, Carb | for *N. benthamiana* expression |
| pL1M-F1-LpCathB::2x35S | GV3101 | Rif, Gent, Carb | for *N. benthamiana* expression |
| pL1M-F1-CP1Amut-nogran-mCherry::2x35S | GV3101 | Rif, Gent, Carb | for *N. benthamiana* expression |
| pL1M-F3_2x35S-p19_pterm | GV3101 | Rif, Gent, Carb | for *N. benthamiana* expression |
| pRSET-PP-LpCys1-noSP | BL21 | Carb, Clm | for *E. coli* expression |

**Supplementary Table S3:** Identifiers used in Byrne *et al.,* (2015) and this study.

| ID in Byrne *et al*., 2015: | short ID: | Name: |
|---|---|---|
| maker-scaffold_4213\|ref0036040-exonerate_est2genome-gene-0.0-mRNA-1 | R_4213 | |
| maker-scaffold_1277\|ref0038680-exonerate_est2genome-gene-0.0-mRNA-1 | R_1277 | |
| maker-scaffold_17473\|ref0028128-exonerate_est2genome-gene-0.0-mRNA-1 | R_17473 | |
| maker-scaffold_13429\|ref0032027-exonerate_est2genome-gene-0.0-mRNA-1 | R_13429.1 | |
| maker-scaffold_13429\|ref0032027-exonerate_est2genome-gene-0.0-mRNA-2 | R_13429.2 | |
| maker-scaffold_943\|ref0030985-exonerate_est2genome-gene-0.0-mRNA-1 | R_943 | |
| maker-scaffold_16198\|ref0011939-exonerate_est2genome-gene-0.0-mRNA-1 | R_16198 | |
| maker-scaffold_1418\|ref0028322-exonerate_est2genome-gene-0.1-mRNA-1 | R_1418 | |
| maker-scaffold_11784\|ref0004048-exonerate_est2genome-gene-0.1-mRNA-1 | R_11784 | |
| maker-scaffold_4437\|ref0034794-exonerate_est2genome-gene-0.2-mRNA-1 | R_4437 | |
| maker-scaffold_4870\|ref0016801-exonerate_est2genome-gene-0.3-mRNA-1 | R_4870 | LpCP2 |
| maker-scaffold_7574\|ref0028846-exonerate_est2genome-gene-0.4-mRNA-1 | R_7574 | |
| maker-scaffold_11332\|ref0028120-exonerate_est2genome-gene-0.0-mRNA-1 | R_11332 | |
| maker-scaffold_2516\|ref0039699-exonerate_est2genome-gene-0.3-mRNA-1 | R_2516 | LpCP1 |
| maker-scaffold_8459\|ref0014698-exonerate_est2genome-gene-0.0-mRNA-1 | R_8459 | |
| maker-scaffold_182\|ref0000331-exonerate_est2genome-gene-0.0-mRNA-1 | R_182 | LpXCP2 |
| maker-scaffold_937\|ref0025736-exonerate_est2genome-gene-0.0-mRNA-1 | R_937 | |
| maker-scaffold_11872\|ref0015306-exonerate_est2genome-gene-0.3-mRNA-1 | R_11872 | LpCathB |
| maker-scaffold_2102\|ref0004996-exonerate_est2genome-gene-0.0-mRNA-1 | R_2102 | |
| maker-scaffold_82\|ref0023171-exonerate_est2genome-gene-1.2-mRNA-1 | R_82 | |
| maker-scaffold_393\|ref0000952-exonerate_est2genome-gene-0.5-mRNA-1 | R_393 | |
| maker-scaffold_3210\|ref0042499-exonerate_est2genome-gene-0.0-mRNA-1 | R_3210 | |
| maker-scaffold_3760\|ref0018795-exonerate_est2genome-gene-0.0-mRNA-1 | R_3760 | |
| maker-scaffold_12141\|ref0031444-exonerate_est2genome-gene-0.0-mRNA-1 | R_12141 | LpCys1 |
| maker-scaffold_2071\|ref0020178-exonerate_est2genome-gene-0.0-mRNA-1 | R_2071 | LpCys4 |
| maker-scaffold_27228\|ref0035705-exonerate_est2genome-gene-0.0-mRNA-1 | R_27228 | LpCys9 |

**Supplementary Table S4:** Identification of PLCP inhibitor candidates in *E. festucae* genome.

| **Inhibitor** | **Pathogen** | **Homologue ID** | **E-value** |
|---|---|---|---|
| Avr2 | *Passalora fulva* | Fl1_007263<br>Fl1_005391 | 2.8<br>5.9 |
| EpiC1 | *Phytophthora infestans* | Fl1_001831<br>Fl1_004526 | 4.0<br>6.2 |
| VAP1 | *Globodera rostochiensis* | Fl1_007387<br>Fl1_004109 | 6e-09 *<br>1e-04 * |
| Avrblb2 | *Phytophthora infestans* | Fl1_004423<br>Fl1_000203 | 0.3<br>1.2 |
| Pit2 | *Ustilago maydis* | Fl1_000888<br>Fl1_003250 | 2.4<br>3.8 |
| popP2 | *Ralstonia* solanacearum | Fl1_005012<br>Fl1_004802 | 0.003<br>0.85 |
| SDE1 | *Candidatus Liberibacter asiaticus* | Fl1_003013<br>Fl1_005446 | 0.5<br>1.0 |
| Cip1 (PSPTO_4211) | *Pseudomonas syringae pv. tomato str. DC3000* | Fl1_002321<br>Fl1_006287 | 1.4<br>2.5 |

The two best hits found for each inhibitor are displayed in this table.
* E-values with significant homology.

**Supplementary Table S5:** functional annotation of Fl1 “unknown” proteins identified in apoplastic fluid of infected *L. perenne* plants.

| ID: | Prediction apoplastP 1.0 | apoplastP Probability | Prediction EffectorP | EffectorP Probability | Quantified in MS |
|---|---|---|---|---|---|
| Fl1_007060 | Non-apoplastic | 0.99 | Non-effector | 1 | Yes |
| Fl1_000097 | Apoplastic | 0.59 | Non-effector | 1 | Yes |
| Fl1_000292 | Non-apoplastic | 1 | Non-effector | 1 | Yes |
| Fl1_000648 | Non-apoplastic | 0.99 | Non-effector | 1 | Yes |
| Fl1_001039 | Non-apoplastic | 0.7 | Effector | 0.71 | Yes |
| Fl1_001817 | Non-apoplastic | 0.96 | Non-effector | 1 | Yes |
| Fl1_001861 | Apoplastic | 0.75 | Non-effector | 1 | Yes |
| Fl1_002355 | Apoplastic | 0.68 | Non-effector | 1 | Yes |
| Fl1_002809 | Non-apoplastic | 0.73 | Non-effector | 0.997 | Yes |
| Fl1_003471 | Apoplastic | 0.77 | Effector | 0.999 | Yes |
| Fl1_003555 | Apoplastic | 0.55 | Non-effector | 0.667 | Yes |
| Fl1_003640 | Non-apoplastic | 0.89 | Non-effector | 1 | Yes |
| Fl1_005600 | Apoplastic | 0.66 | Non-effector | 0.834 | Yes |
| Fl1_006178 | Non-apoplastic | 0.92 | Non-effector | 1 | Yes |
| Fl1_006292 | Apoplastic | 0.66 | Non-effector | 1 | Yes |
| Fl1_006293 | Apoplastic | 0.84 | Non-effector | 1 | Yes |
| Fl1_006865 | Non-apoplastic | 0.8 | Non-effector | 1 | Yes |
| Fl1_007049 | Non-apoplastic | 0.76 | Effector | 0.89 | Yes |
| Fl1_007055 | Non-apoplastic | 0.53 | Non-effector | 1 | Yes |
| Fl1_007463 | Non-apoplastic | 0.51 | Non-effector | 1 | Yes |
| Fl1_001500 | Non-apoplastic | 0.62 | Effector | 0.974 | No |
| Fl1_001801 | Apoplastic | 0.93 | Non-effector | 1.0 | No |
| Fl1_002618 | Non-apoplastic | 0.63 | Non-effector | 1.0 | No |
| Fl1_002869 | Apoplastic | 0.84 | Effector | 0.995 | No |
| Fl1_003333 | Apoplastic | 0.78 | Effector | 0.997 | No |
| Fl1_003341 | Apoplastic | 0.71 | Non-effector | 1.0 | No |
| Fl1_004066 | Non-apoplastic | 0.73 | Non-effector | 1.0 | No |
| Fl1_004767 | Apoplastic | 0.61 | Non-effector | 1.0 | No |
| Fl1_004793 | Apoplastic | 0.79 | Non-effector | 0.999 | No |
| Fl1_005030 | Non-apoplastic | 0.77 | Effector | 0.578 | No |
| Fl1_005116 | Apoplastic | 0.75 | Non-effector | 0.86 | No |
| Fl1_005240 | Apoplastic | 0.83 | Effector | 0.972 | No |
| Fl1_005423 | Non-apoplastic | 0.99 | Non-effector | 1.0 | No |
| Fl1_006033 | Apoplastic | 0.8 | Non-effector | 1.0 | No |
| Fl1_006202 | Apoplastic | 0.72 | Effector | 0.874 | No |
| Fl1_006971 | Non-apoplastic | 0.99 | Non-effector | 1.0 | No |
| Fl1_007061 | Non-apoplastic | 0.68 | Non-effector | 1.0 | No |

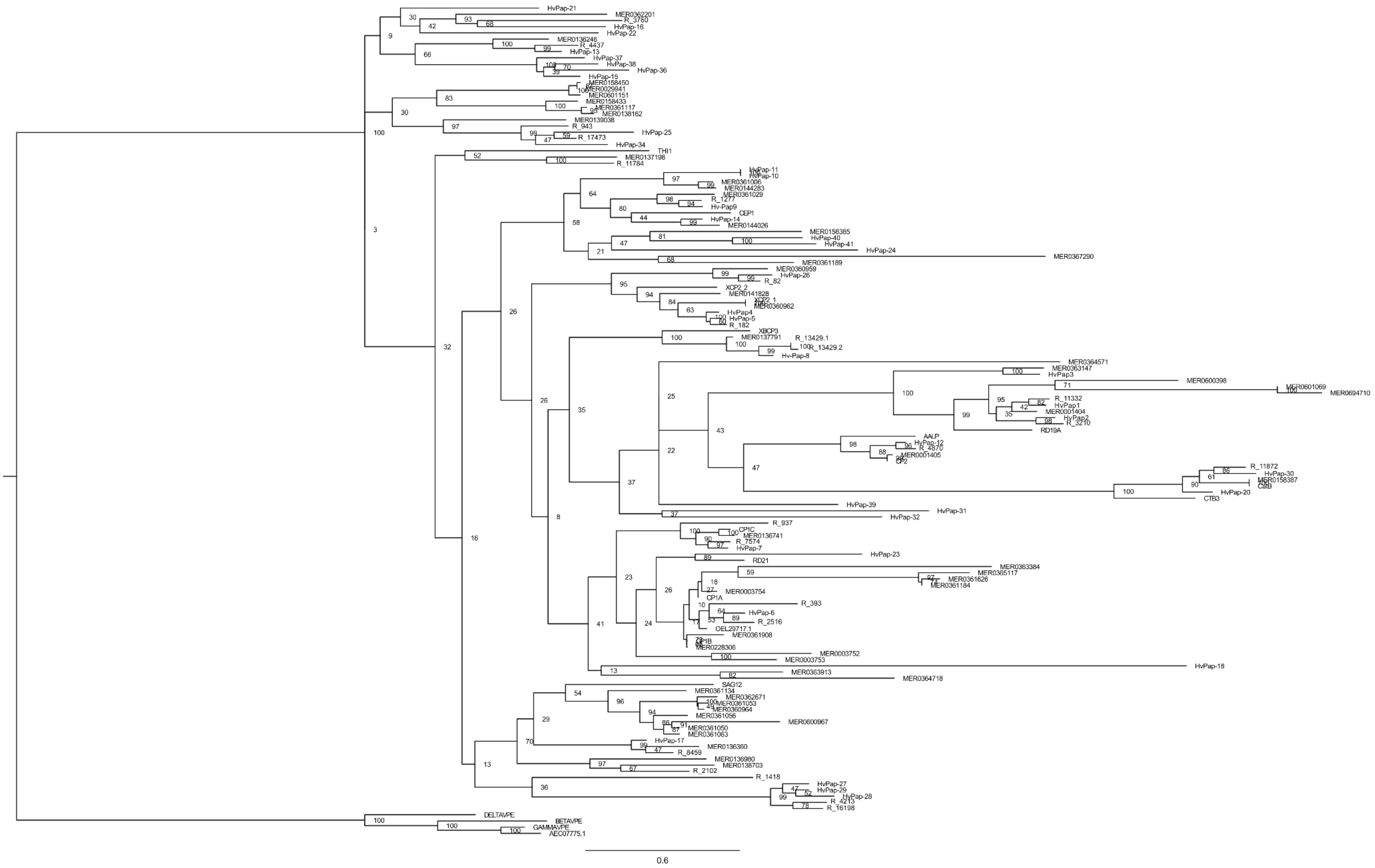

**Supplementary Fig. S1.** Unedited phylogenetic tree of *L. perenne* PLCPs. Phylogenetic analysis was performed using 23 *L. perenne* PLCP sequences identified via functional domain analysis (R numbers), 52 PLCP sequences from the maize line B73 obtained from MEROPS database (www.ebi.ac.uk/merops), 6 PLCP sequences from the maize line EGB, and 38 PLCP sequences of *Hordeum vulgare* (HvPaps). The four legumains (AtVPEs) from *A. thaliana* were used to root the phylogenetic three and one member of each PLCP subfamily also from *A. thaliana* for the subfamily classification. In this analysis, full length protein sequences were used, including signal peptide, auto-inhibitory pro-domain, protease C1-domain and if present granulin domain.

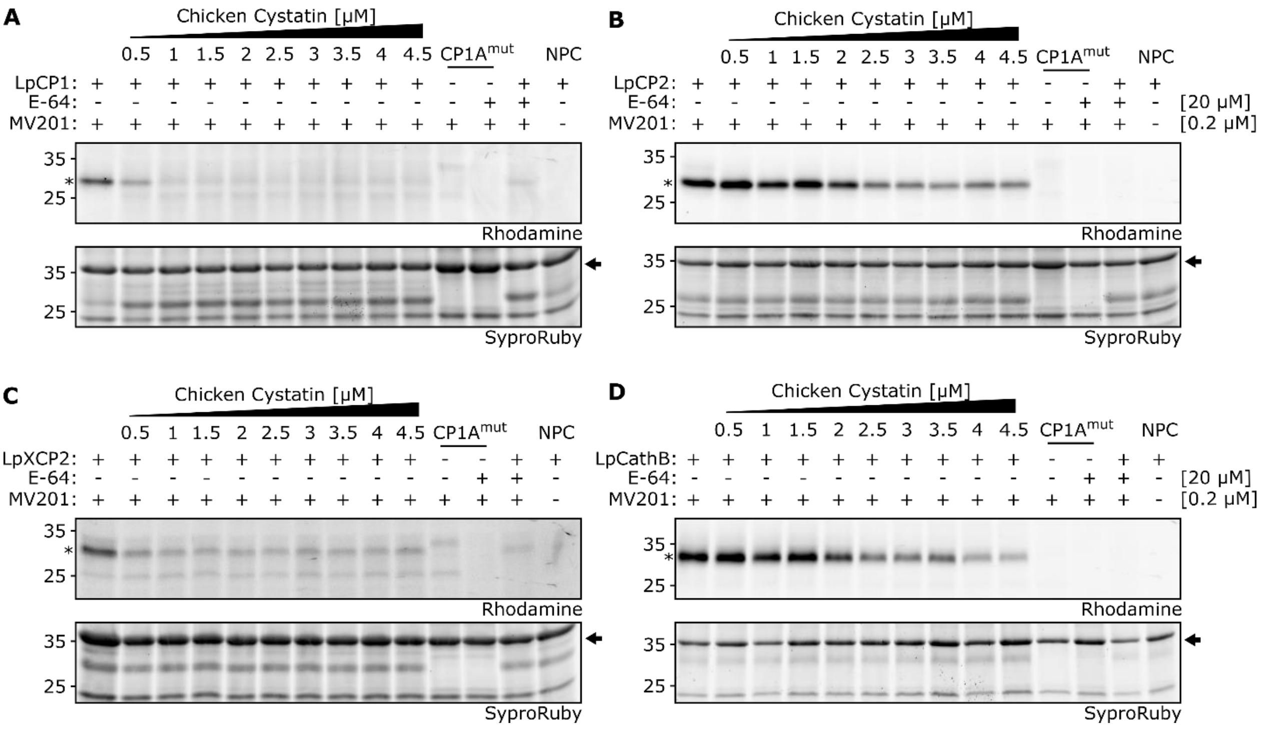


**Supplementary Fig. S2.** Inhibitory activity of CWEC on overexpressed ryegrass PLCPs. Four PLCPs of *L. perenne* were overexpressed in *N. benthamiana* using Agrobacterium-mediated transformation. Apoplastic fluid containing PLCPs was isolated and monitored using the fluorescent probe MV201. Samples were pre-incubated for 15 min with 20 µM E-64 or a concentration range (0 – 4.5 µM) of CWEC followed by 2 h labelling with 0.2 µM MV201. As background controls, samples containing CP1A$^{mut}$, an inactive maize PLCP, and the no-probe control (NPC) were prepared. The activity and inhibitory effect of CWEC on LpCP1 (A), LpCP2 (B), LpXCP2 (C) and LpCathB (D) was analyzed using in gel fluorescent scanning and as a loading control SyproRuby staining was performed. For each analyzed PLCP (marked as asterisk) signals were quantified and normalized to a loading control signal (marked with an arrow) and activity without inhibitor was set to 100% (Fig. 7).
